# ERBB2 establishes the timing of progesterone priming required for uterine receptivity

**DOI:** 10.64898/2026.09.14.751518

**Authors:** Bo Li, Chen Zhang, Amanda Dewar, Xiaoli Liu, Wenbo Deng, Hongbo Qi, Sudhansu K. Dey, Xiaofei Sun

## Abstract

Progesterone is essential for establishing uterine receptivity and sustaining early pregnancy. However, how the uterus acquires the competence to respond to progesterone at the appropriate time remains poorly understood. Here we show that ERBB2 governs the timing of progesterone responsiveness required for uterine receptivity. In endometrial samples from women with recurrent spontaneous abortion (RSA), ERBB2 is downregulated specifically in the stromal compartment, accompanied by a parallel reduction in stromal progesterone receptor (PGR). Uterine deletion of *Erbb2* in mice blunts the rise of stromal PGR at the onset of the receptive window, deferring implantation and impairing decidualization despite normal circulating progesterone and estrogen. Epithelial deletion has no such effect, placing the requirement in the stroma. High-dose progesterone fails to rescue the phenotype; advancing progesterone priming by 24 hours restores stromal PGR, HAND2, *Bmp2*, and on-time implantation. Collectively, the present study provides evidence that ERBB2 directs the temporal window of stromal progesterone priming by setting the threshold for PGR induction, thereby synchronizing the maternal-fetal interface and safeguarding pregnancy.

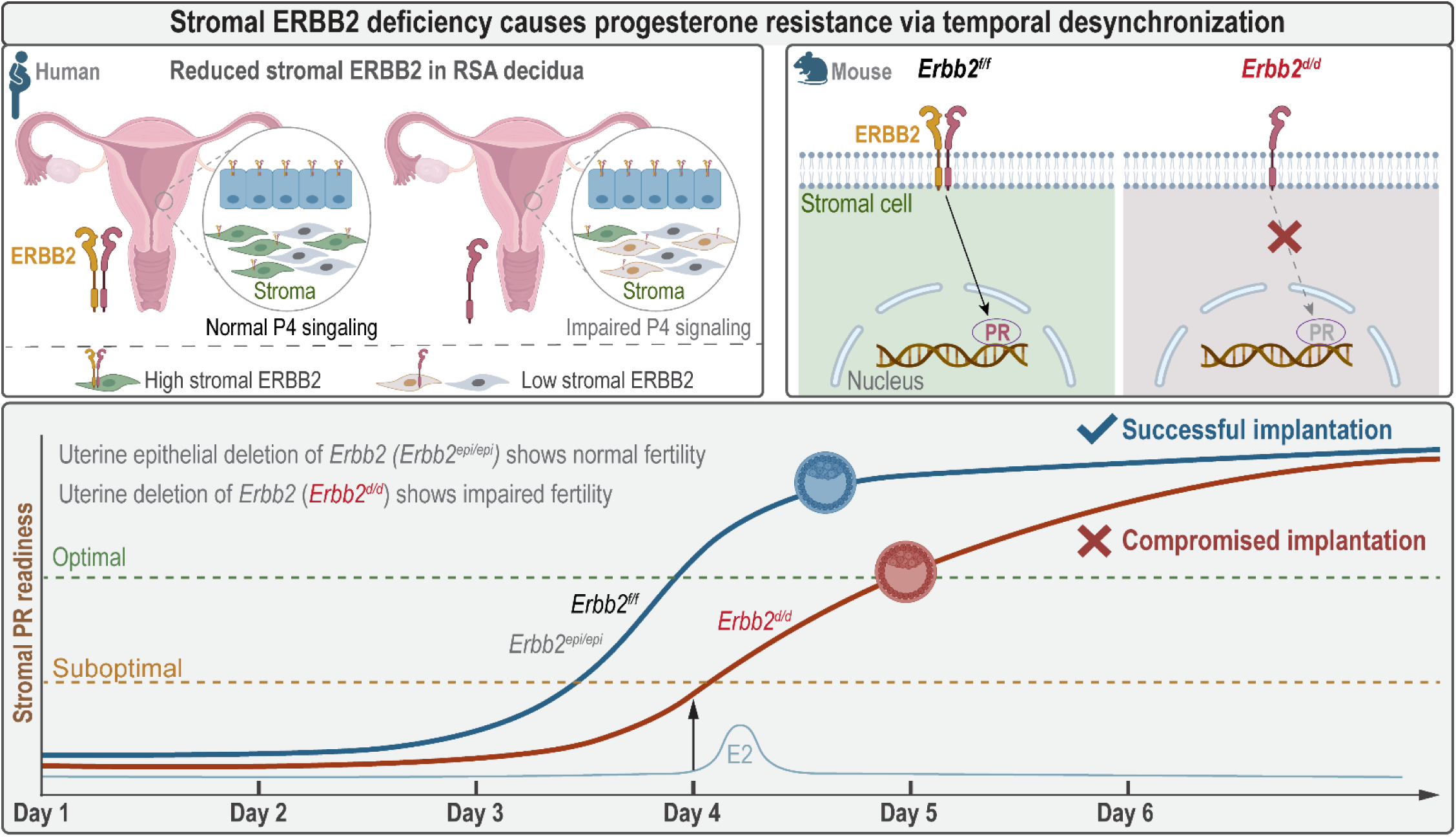

## Introduction

Recurrent spontaneous abortion (RSA), defined as the loss of two or more consecutive pregnancies, affects 1– 2% of reproductive-age couples (1). Although chromosomal aneuploidy accounts for a portion of sporadic losses, more than half of RSA cases remain unexplained after standard clinical evaluation (2). Systemic progesterone supplementation is widely used in management, yet a substantial subset of patients fails to respond despite normal circulating hormone levels, a condition termed “progesterone resistance.” The implantation failure and defective decidualization observed in these patients indicate that the principal defect resides not in the endocrine signal itself but in the intrinsic capacity of the endometrial stroma to mount a progesterone response. How the stroma acquires this competence, and what molecular mechanism sets its timing, remains unknown. Single-cell transcriptomic analysis of RSA decidua points toward the answer: the stromal compartment in these patients is marked by depleted functional decidual subpopulations, accumulated senescence-associated cells, and aberrant differentiation trajectories, a “defective stromal niche” whose molecular basis has not been resolved (3).

Successful pregnancy hinges on decidualization, a profound tissue remodeling event where elongated stromal fibroblasts differentiate into specialized, secretory epithelioid cells (4). Crucially, this cellular transformation is principally driven by progesterone signaling mediated through the nuclear progesterone receptor (5). While fundamental to placental mammals, the initiation of this process exhibits distinct species-specific strategies. In humans, decidualization is spontaneous: initiated by rising post-ovulatory progesterone levels, the stroma begins to differentiate and create a receptive environment before the embryo even arrives. Conversely, murine decidualization is induced (4). In the mouse, progesterone signaling first regulates downstream targets, such as *Ihh* and *Hoxa10*, to establish uterine receptivity (6). Only after the blastocyst successfully attaches (implantation) does the decidual program physically begin (7). Despite this temporal reversal, “preparation” in humans versus “reaction” in mice, the downstream molecular machinery governing stromal differentiation remains highly conserved and strictly progesterone-dependent in both species. Thus, defects in this shared intrinsic program disrupt the maternal-fetal dialogue, rendering the endometrium hostile to implantation and driving RSA (1).

The epidermal growth factor (EGF) family, comprising EGF, transforming growth factor-α (TGF-α), heparin-binding EGF (HBEGF), amphiregulin (Ar), betacellulin, epiregulin, and heregulins/neu differentiation factors (NDFs), plays a critical role in various biological processes (8, 9). These ligands are synthesized as transmembrane proteins and undergo proteolytic cleavage to release their active forms. They exert their effects through interaction with the ERBB receptor family, which includes four receptor tyrosine kinases: ERBB1 (EGFR), ERBB2, ERBB3, and ERBB4 (10). While these receptors share structural similarities, they exhibit distinct ligand-binding specificities and kinase activities, contributing to the diverse signaling outcomes mediated by the EGF family. ERBB2, lacking a direct ligand, has the strongest kinase activity and preferentially dimerizes with other ERBB family members. In contrast, ERBB3, with impaired kinase activity, requires heterodimerization for intracellular signaling (11). HBEGF is an early marker mediating embryo-uterine crosstalk. It directly binds to EGFR and ErbB4 to facilitate embryo growth (12, 13). Our previous studies demonstrate that HBEGF promotes Vangl2 phosphorylation via ERBB2/ERBB3 heterodimers during embryo implantation (14). Based on our previous findings, *Erbb2* mRNA expression expands from epithelial cells during the peri-implantation stage (days 1-4) to stromal cells within the primary decidual zone (PDZ) and secondary decidual zone (SDZ), as demonstrated by *in situ* hybridization (15). Although our lab has confirmed that uterine specific deletion of *Erbb2* (*Erbb2^f/f^ Pgr^Cre/+^*, also named as *Erbb2^d/d^*) leads to reduced litter sizes, and uterine epithelial deletion of *Erbb2* (*Erbb2^f/f^ Ltf^Cre/+^*, also named as *Erbb2^epi/epi^*) shows normal pregnancy (14), the underlying mechanisms by which *Erbb2* functions during early pregnancy remain unclear.

Here, we bridge clinical transcriptomics with mechanistic genetics to define the hierarchy of stromal regulation. Through re-analysis of single-cell RNA sequencing data from patients with RSA, we identify a downregulation of *ERBB2* in the stromal compartment. Using a uterine-specific knockout mouse model, we further demonstrate that ERBB2 acts as an essential temporal gatekeeper of stromal progesterone receptor (PGR) activation. We report that loss of *Erbb2* blunts the initial induction of stromal PGR at the onset of receptivity, resulting in deferred implantation and impaired decidualization despite normal systemic hormone levels. Critically, we show that this defect is time-dependent, as optimizing the timing of progesterone exposure rescues the phenotype. These findings establish stromal *ERBB2* deficiency as a driver of “progesterone-resistant” pregnancy loss and highlight a previously unrecognized molecular timer essential for synchronizing the maternal-fetal interface.

## Results

### ERBB family receptor deficiency is associated with human recurrent spontaneous abortion

To ask whether endometrial defects underlie early pregnancy loss, we reanalyzed a public single-cell RNA-seq dataset of 66,078 cells from women with RSA and healthy controls. t-SNE clustering resolved the major uterine compartments (Figure 1A to C) and allowed us to examine ERBB receptor expression in each.

**Figure 1.**
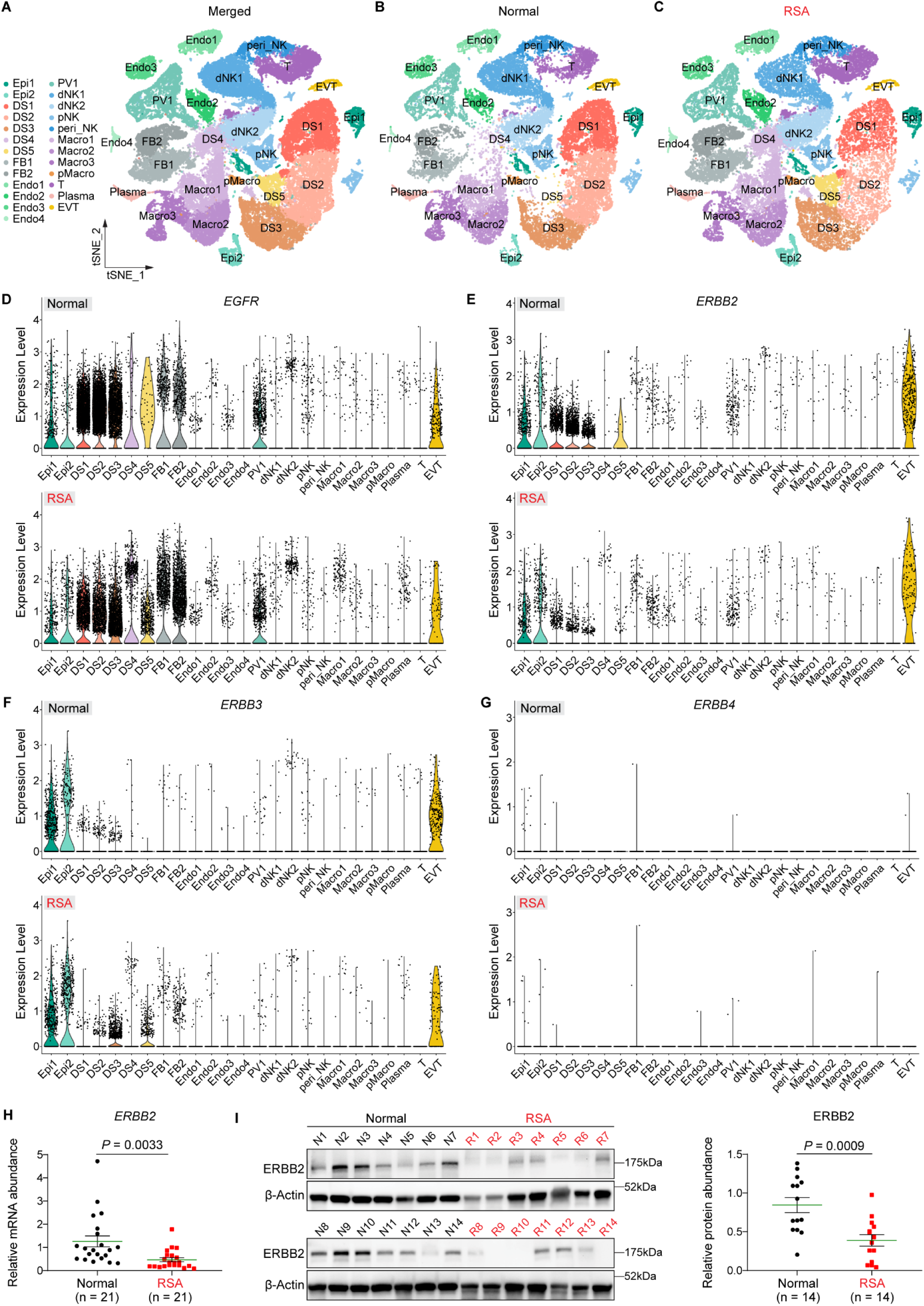
ERBB family receptor deficiency characterizes human recurrent spontaneous abortion. (**A** to **C**) t-SNE visualization of single-cell transcriptomes from decidual tissues of healthy controls (Normal) and patients with recurrent spontaneous abortion (RSA). (**A**) Merged projection of 66,078 cells revealing major cell populations. (**B** and **C**) Cellular distribution patterns separated by two groups: Normal (**B**) and RSA (**C**). (**D** to **G**) Violin plots comparing the expression profiles of ERBB family receptor tyrosine kinases, *EGFR* (**D**), *ERBB2* (**E**), *ERBB3* (**F**), and *ERBB4* (**G**), across cell clusters in Normal versus RSA tissues. (**H**) qPCR validation of *ERBB2* mRNA abundance in an independent clinical cohort of Normal (n = 21) and RSA (n = 21) endometrial tissues. (**I**) Representative Western blot and quantification of ERBB2 protein levels in matched tissue samples. Data in (**H**) and (**I**) are mean ± SEM. *P* values were determined by unpaired two-tailed Student’s *t*-test.

EGFR and ERBB2 were unchanged in the epithelium (Figure 1D and E) but both were downregulated in the RSA stroma. ERBB3 and ERBB4 showed no difference in either compartment (Figure 1F and G). EGFR is a known regulator of implantation and decidualization (16), but ERBB2 has no defined function in the uterine stroma. Its loss in RSA endometrium was therefore unexpected, and we focused on stromal ERBB2.

To validate this finding, we collected endometrial tissue from an independent cohort of women with RSA (n = 21) and fertile controls (n = 21). *ERBB2* mRNA was reduced in RSA endometria by qPCR (*P* = 0.0033; Figure 1H), and ERBB2 protein was correspondingly lost by Western blot (P = 0.0009; Figure 1I). Together with the single-cell data, these results identify stromal ERBB2 deficiency as a reproducible feature of RSA.

### Uterine deletion of *Erbb2* compromises pregnancy outcomes

The consistent loss of stromal ERBB2 in RSA raised the question of whether uterine *Erbb2* is required for normal pregnancy progression. To address this, we generated uterine-specific *Erbb2* knockout mice (*Erbb2^d/d^*) and examined pregnancy outcomes. Although *Erbb2^d/d^* females showed normal pregnancy rates and pup weights, litter sizes were consistently reduced compared with *Erbb2^f/f^* controls (Figure 2A to C). Tracing back from delivery, *Erbb2^d/d^* uteri on day 16 contained fewer implantation sites, and by day 12 the rate of resorption sites was elevated, and gross implantation site weights were significantly decreased (Figure 2D and E). On day 8, implantation site number was comparable between genotypes, but decidual weights were markedly reduced in *Erbb2^d/d^*uteri (Figure 2F). Histological analysis revealed that embryos within *Erbb2^d/d^* uteri were compromised and the secondary decidual zone was substantially smaller than in floxed controls (Figure 2F). Taken together, these results demonstrate that uterine deletion of *Erbb2* impairs decidualization and pregnancy outcomes without affecting overall implantation frequency.

**Figure 2.**
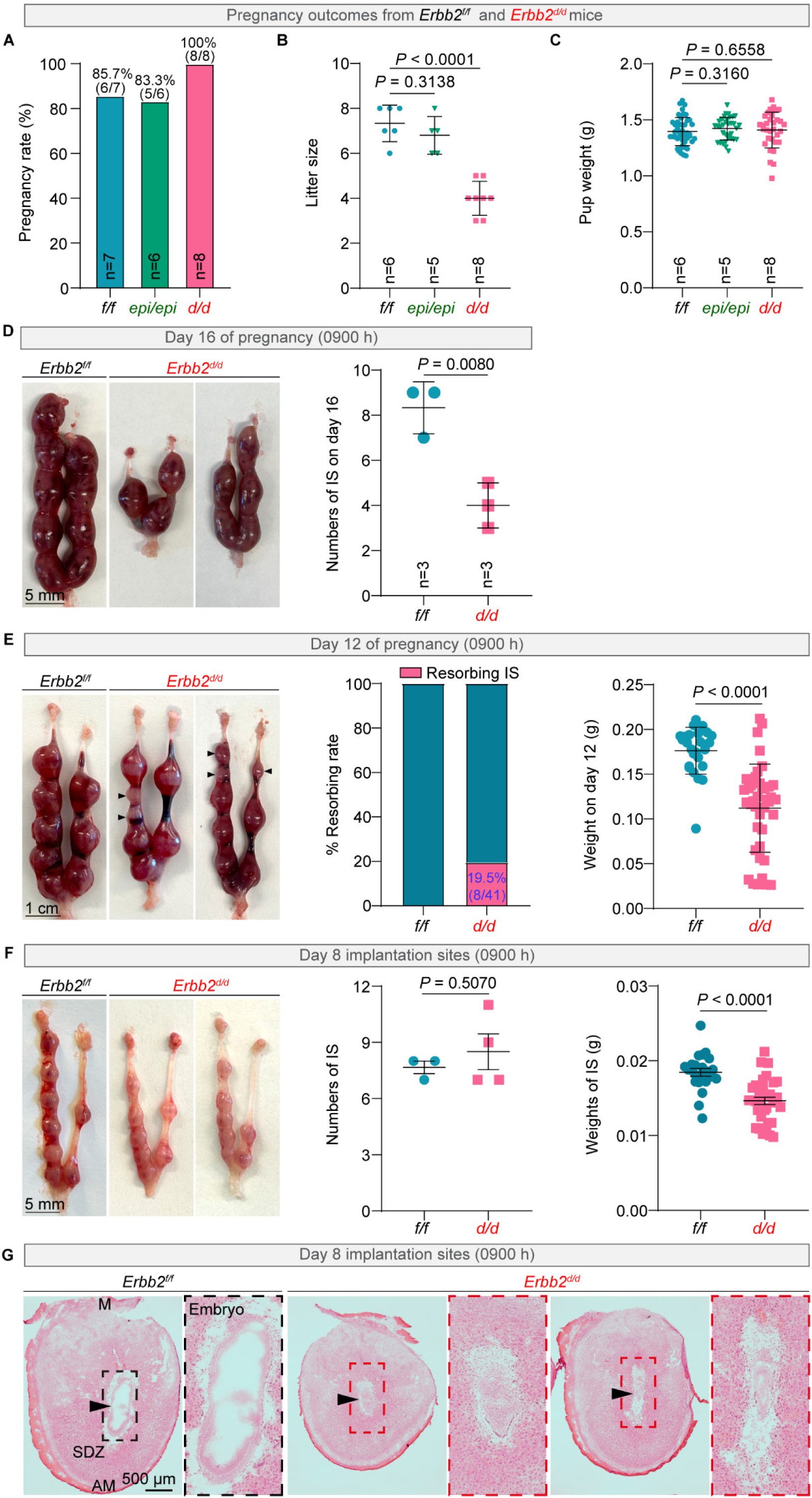
Uterine deletion of *Erbb2* reduces litter sizes. (**A**) Pregnancy rate (*f/f*: n = 7; *epi/epi*: n = 6; *d/d*: n = 7) in *Erbb2^f/f^*, *Erbb2^epi/epi^*, *and Erbb2^d/d^* mice. Data are presented as mean ± SEM. (**B**) Litter size (*f/f*: n = 6; *epi/epi*: n = 5; *d/d*: n = 8) in *Erbb2^f/f^*, *Erbb2^epi/epi^*, *and Erbb2^d/d^* mice. Data are presented as mean ± SEM. (**C**) Pup weights (*f/f*: n = 6; *epi/epi*: n = 5; *d/d*: n = 8) in *Erbb2^f/f^*, *Erbb2^epi/epi^*, *and Erbb2^d/d^* mice. Data are presented as mean ± SEM. (**D**) Representative images of uteri collected from *Erbb2^f/f^* (n = 3) and *Erbb2^d/d^* (n = 3) females on day 16 of pregnancy at 0900 h, along with the number of implantation sites (IS) on day 16. Scale bar: 5 mm. Quantified data (*f/f*: n = 3; *d/d*: n = 3) are presented as mean ± SEM. (**E**) Representative images of day 12 uteri in *Erbb2^f/f^* (n = 3) and *Erbb2^d/d^* (n = 5) females. Scale bar: 1 cm. Percentage of resorbing implantation sites (% Resorbing IS) on day 12 of pregnancy in *Erbb2^f/f^* (n = 3) and *Erbb2^d/d^* (n = 5) females. Embryo weights on day 12 of pregnancy. Each dot represents one implantation site. Data are presented as mean ± SEM. (**F**) Representative images of day 8 pregnant uteri from *Erbb2^f/f^*(n = 3) and *Erbb2^d/d^* (n = 4) females, showing the number of implantation sites and related decidual weights. Scale bar: 5 mm. Data are presented as mean ± SEM. **(G)** Histology of day 8 implantation sites from *Erbb2^f/f^* (n = 3) and *Erbb2^d/d^* (n = 3) females. Arrowheads indicate the location of embryos. M mesometrial pole, AM antimesometrial pole. Scale bar: 500 μm.

### Uterine deletion of *Erbb2* impairs decidualization

The reduction in decidual weight prompted us to ask whether the decidualization program itself is compromised in *Erbb2^d/d^* uteri. We examined bone morphogenetic protein 2 (*Bmp2*), a well-established marker of decidualization (17). FISH results show that *Bmp2* is predominantly expressed in stromal cells surrounding the implanting blastocyst on day 5, contributing to the formation of the primary decidual zone (PDZ). However, *Erbb2^d/d^* uteri showed reduced *Bmp2* expression (Figure 3A), indicating an impairment in the initial decidualization process following uterine deletion of *Erbb2*. As decidualization progresses, *Bmp2* expands its expression from primary decidual zone to the secondary decidual zone (SDZ) from day 5 to day 6 (18, 19). As expected, we also observed reduced *Bmp2* expression in the SDZ on day 6 in *Erbb2^d/d^* uteri (Figure 3B). Heart and neural crest derivatives-expressed protein 2 (HAND2), a progesterone-induced transcription factor, is specifically expressed in the uterine stroma and is essential for embryo implantation and decidualization (20). Immunostaining shows that HAND2 is mainly localized in the primary decidual zone of *Erbb2^f/f^* uterus but is substantially decreased in *Erbb2^d/d^* uteri on day 5 (Figure 3C). As HAND2 is also known for its anti-proliferative function, we observed intense cell proliferation marked by Ki67 staining in stromal cells surrounding the implanting blastocysts in *Erbb2^d/d^* uteri. In contrast, stromal cells surrounding the embryo in the floxed mice show halted cell proliferation (Figure 3D). These findings demonstrate that uterine deletion of *Erbb2* impairs stromal cell decidualization. To determine whether *Erbb2* is solely required for decidualization, we utilized two in vitro decidualization markers, *Prl8a2* and *Prl3c1*, to assess. Our results demonstrated that mRNA levels of *Prl8a2* and *Prl3c1* were significantly decreased in *Erbb2^d/d^* uterine stromal cells after 3 or 6 days of in vitro decidualization compared to the *Erbb2^f/f^* control group (Figure 3E and F). Collectively, these results suggest that uterine deletion of *Erbb2* impairs mouse decidualization.

**Figure 3.**
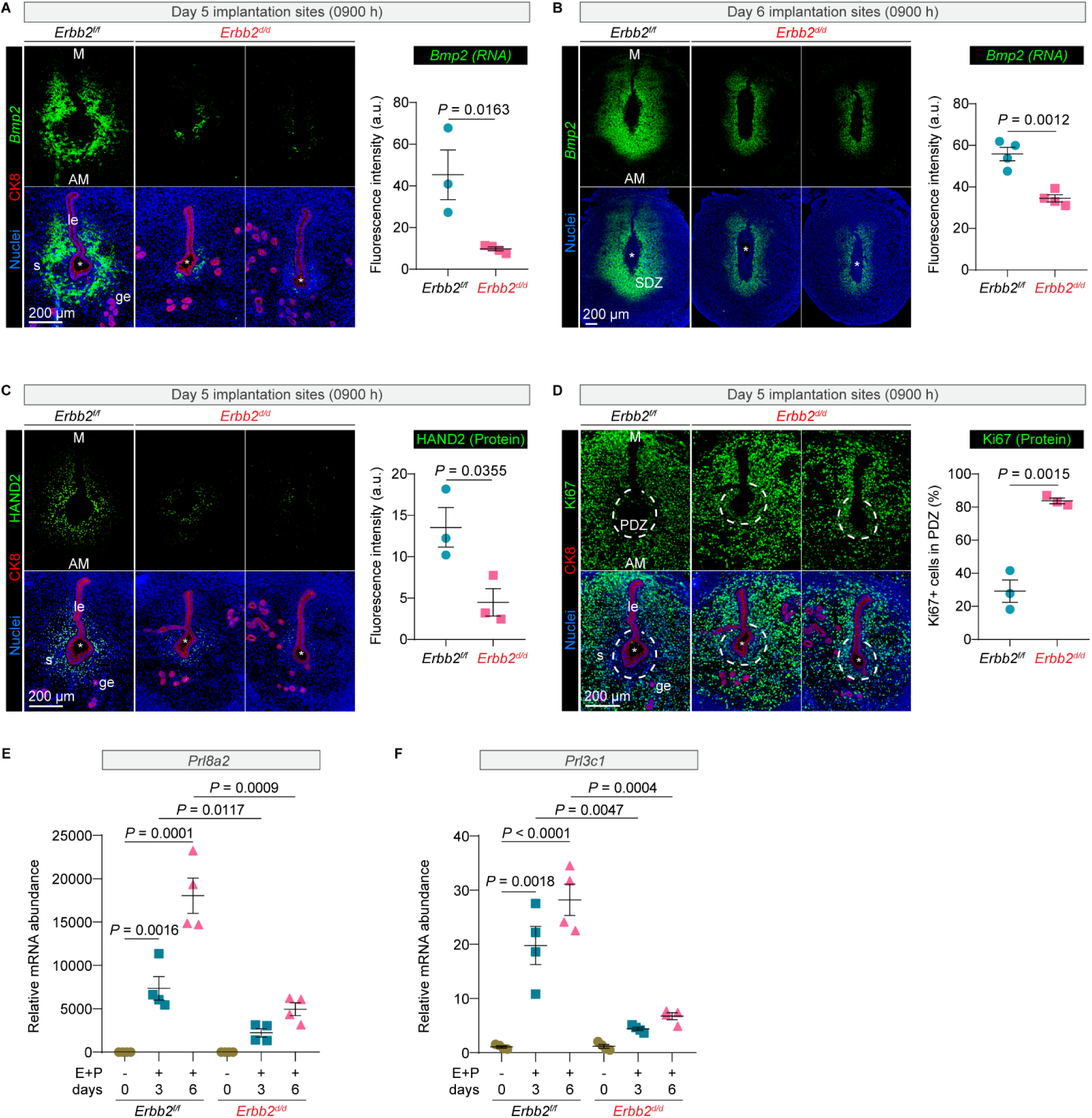
*Erbb2* deletion impairs stromal cell decidualization. (**A**) Representative images of *Bmp2 mRNA* detected by FISH and CK8 immunofluorescence in day 5 implantation sites (0900 h) from *Erbb2^f/f^*(n = 3) and *Erbb2^d/d^* (n = 4) females. Nuclei are counterstained with DAPI. Quantification of *Bmp2* fluorescence intensity is shown on the right. (**B**) FISH analysis of *Bmp2* mRNA in day 6 implantation sites (0900 h) from *Erbb2^f/f^* (n = 4) and *Erbb2^d/d^* (n = 4) females, with quantification of fluorescence intensity. (**C**) Immunofluorescence of HAND2 and CK8 in day 5 implantation sites (0900 h) from *Erbb2^f/f^* (n = 3) and *Erbb2^d/d^* (n = 3) females, with quantification of HAND2 fluorescence intensity. Scale bar: 200 μm. (**D**) Immunofluorescence of Ki67 and CK8 in day 5 implantation sites (0900 h) from *Erbb2^f/f^* (n = 3) and *Erbb2^d/d^* (n = 3) females. Dashed lines indicate the primary decidual zone (PDZ). Scale bar: 200 μm. Quantification of Ki67⁺ cells within the PDZ are shown on the right. (**E** and **F**) Relative mRNA levels of *Prl8a2* and *Prl3c1* in isolated *Erbb2^f/f^* (n = 4) and *Erbb2^d/d^* (n = 4) stromal cells after 3 and 6 days in vitro decidualization with E_2_ and P_4_. Data are presented as mean ± SEM. Asterisks indicate the location of the embryo. le luminal epithelium, ge glandular epithelium, s stroma, M, mesometrial pole, AM, antimesometrial pole, SDZ secondary decidual zone.

### Loss of *Erbb2* disrupts gland-crypt assembly at implantation sites

Given the compromised decidualization in *Erbb2^d/d^* mice, we next asked whether embryo implantation is affected. On day 5 of pregnancy (day 1 = vaginal plug), *Erbb2^d/d^* mice display attenuated blue band responses compared with *Erbb2^f/f^* controls (Figure 4A), a pattern that persists on day 6 (Figure 4B). The total number of implantation sites is comparable between genotypes (Figure 4A and B), indicating that the defect is specific to implantation quality rather than blastocyst attachment frequency. To investigate crypt formation and gland-crypt assembly, we employed 3D imaging after tissue fixation and clearing, as described (21). On day 5, the luminal epithelium invaginates into the decidual stroma to form crypts that house the implanting blastocyst (22); branched glands simultaneously elongate toward the antimesometrial pole, establishing the gland-crypt assembly (21). In *Erbb2^f/f^* uteri, this spatiotemporal program proceeds with well-formed crypts on day 5 (Figure 4C). In *Erbb2^d/d^* uteri, epithelial invagination is absent on day 5, and although partial invagination emerges by day 6, glandular defects persist at implantation sites (Figure 4D). Taken together, these findings demonstrate that *Erbb2* governs gland-crypt assembly during the window of receptivity.

**Figure 4.**
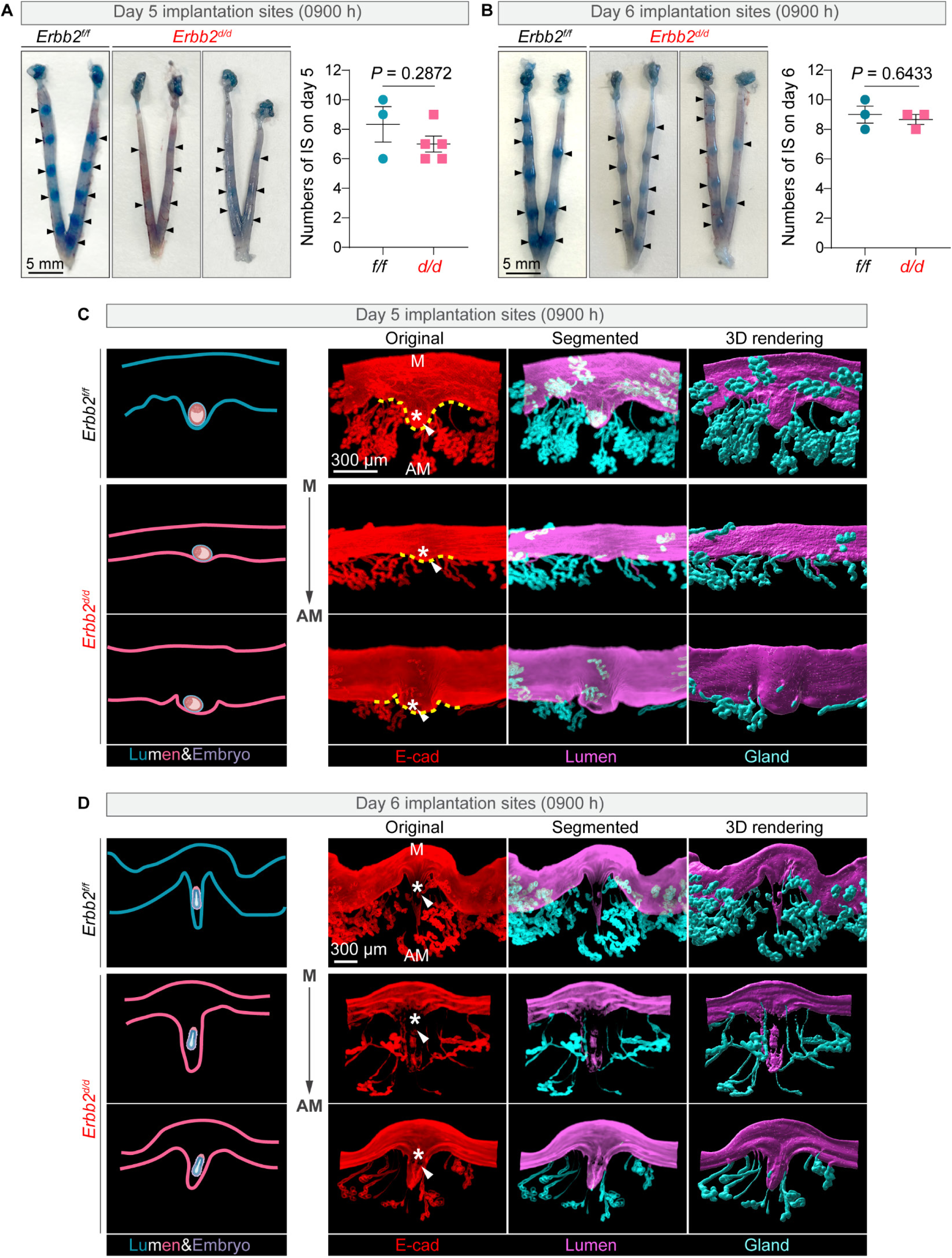
Uterine deletion of *Erbb2* disrupts gland-crypt assembly. (**A** and **B**) Representative images and quantification of uterine implantation sites in *Erbb2^f/f^*and *Erbb2^d/d^* mice on day 5 (**A**; n = 3 for *Erbb2^f/f^*, n = 5 for *Erbb2^d/d^*) and day 6 (**B**; n = 4 for *Erbb2^f/f^*, n = 4 for *Erbb2^d/d^*) of pregnancy. Black arrowheads indicate implantation sites. Data are presented as mean ± SEM. *P* values were determined by unpaired two-tailed Student’s *t*-test. Scale bars: 5 mm. (**C** and **D**) 3D imaging of implantation sites on day 5 (**C**; n = 3 for *Erbb2^f/f^*, n = 5 for *Erbb2^d/d^*) and day 6 (**D**; n = 3 for *Erbb2^f/f^*, n = 3 for *Erbb2^d/d^*). Uterine epithelium was marked by E-cadherin. Scale bars: 300 μm. Yellow dashed lines outline the epithelial crypt. White asterisks indicate the location of the embryo. White arrowheads indicate implantation chambers. M, mesometrial pole; AM, antimesometrial pole.

### Uterine deletion of *Erbb2* defers embryo implantation

In mice, the uterus becomes receptive on day 4 of pregnancy, with the attachment reaction initiating around midnight on the same day (23). Blue bands are evident in *Erbb2^f/f^* mice at midnight on day 4 (referred to as day 4.5). In contrast, *Erbb2^d/d^* uteri showed no blue bands, and blastocysts were recovered by flushing from the *Erbb2^d/d^* uteri (Figure 5A). Heparin-binding EGF-like growth factor (HB-EGF) is the earliest known marker mediating embryo-uterine crosstalk (24, 25). FISH analysis shows that *Hbegf* is expressed in the luminal epithelium and stroma surrounding the implanting blastocyst in *Erbb2^f/f^* uteri but is absent in *Erbb2^d/d^* uteri on day 4 midnight (Figure 5B). Previous studies show that *Cox2* knockout mice exhibit defects in both implantation and decidualization (26). Blastocyst-sized beads pre-coated with HB-EGF elicit responses comparable to those induced by living blastocysts, including the induction of *COX2* (12, 27). COX2 is expressed in the luminal epithelium and stroma surrounding the embryos of *Erbb2^f/f^* uteri on day 4 midnight but is undetectable in *Erbb2^d/d^*uteri. Blastocysts in *Erbb2^d/d^* uteri remain trapped within the luminal epithelium (Figure 5C and D). Although COX2 is absent on day 4.5 in *Erbb2^d/d^* uteri, we found that *Ptgs2*, which encodes COX2, is comparable in *Erbb2^f/f^*and *Erbb2^d/d^* uteri on day 5 of pregnancy (Figure 5E). The uterine epithelium in *Erbb2^d/d^*mice remains intact on day 6, as shown by E-cadherin and β-catenin staining (Figure 5F). Taken together, these findings demonstrate that uterine deletion of *Erbb2* defers embryo implantation.

**Figure 5.**
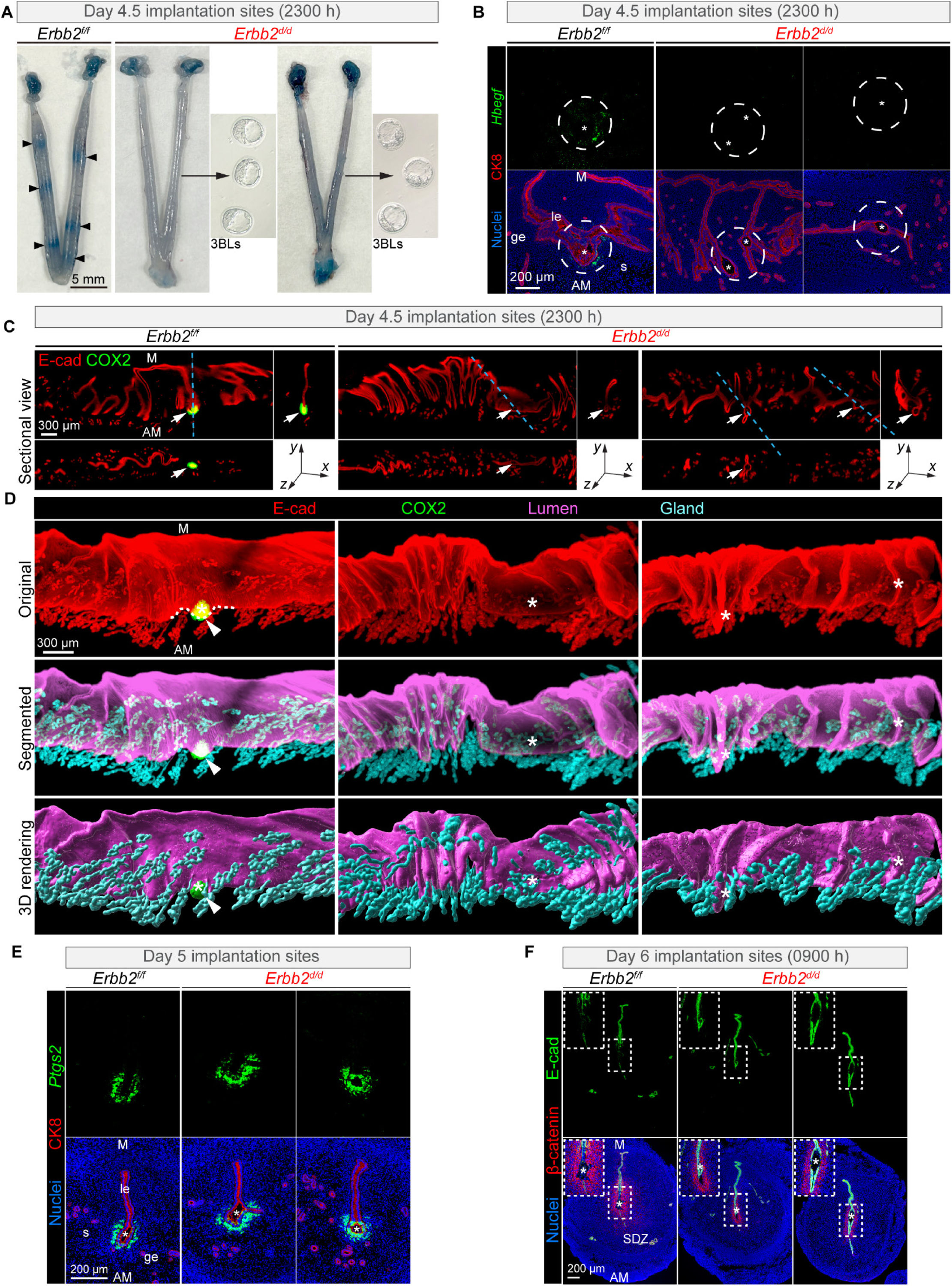
*Erbb2* deletion defers embryo implantation. (**A**) Implantation sites from *Erbb2^f/f^* (n = 3) and *Erbb2^d/d^*(n = 4) uteri at day 4.5 of pregnancy (2300 h). Blastocysts were recovered by flushing uterine horns from the *Erbb2^d/d^* uteri without blue bands. Scale bars, 5 mm. Arrowhead indicates an implantation site. (**B**) FISH of *Hbegf* and Immunofluorescence of CK8 in day 4.5 implantation sites from *Erbb2^f/f^* (n = 3) and *Erbb2^d/d^* (n = 3) uteri. Scale bar, 200 μm. (**C**) 3D imaging of implantation sites at day 4.5 in sectional views. Original (E-cadherin and COX2). Scale bar, 300 μm. (**D**) 3D imaging of implantation sites from *Erbb2^f/f^* and *Erbb2^d/d^* uteri at day 4.5 of pregnancy (2300 h). Scale bar, 300 μm. Original (E-cadherin and COX2), Segmented, and 3D rendering. (**E**) FISH of *Ptgs2* and Immunofluorescence of CK8 on day 5 implantation sites from *Erbb2^f/f^* (n = 3) and *Erbb2^d/d^* (n = 3) uteri. Scale bar, 200 μm. (**F**) Immunofluorescence of E-cadherin and β-catenin in day 6 implantation sites from *Erbb2^f/f^* (n = 5) and *Erbb2^d/d^* (n = 5) uteri. Scale bar, 200 μm. Asterisks indicate the location of the embryo. Arrowheads indicate the implantation chamber (Crypt). le luminal epithelium, ge glandular epithelium, s stroma, M mesometrial pole, AM antimesometrial pole, SDZ secondary decidual zone.

### Uterine deletion of *Erbb2* reduces stromal but not epithelial PGR expression

Although uterine deletion of *Erbb2* defers implantation, embryos in *Erbb2^d/d^* mice still develop to the blastocyst stage (Figure 6A), placing the defect in the uterine compartment rather than the embryo. Given the critical roles of estrogen and progesterone in uterine pregnancy events (28, 29), we measured their serum levels and found no differences between *Erbb2^f/f^* and *Erbb2^d/d^* mice on day 4 (Figure 6B and C). ERα expression is comparable between *Erbb2^f/f^* and *Erbb2^d/d^* uteri on day 4 (Figure 6D), indicating that the defect is not attributable to altered ovarian hormone signaling.

**Figure 6.**
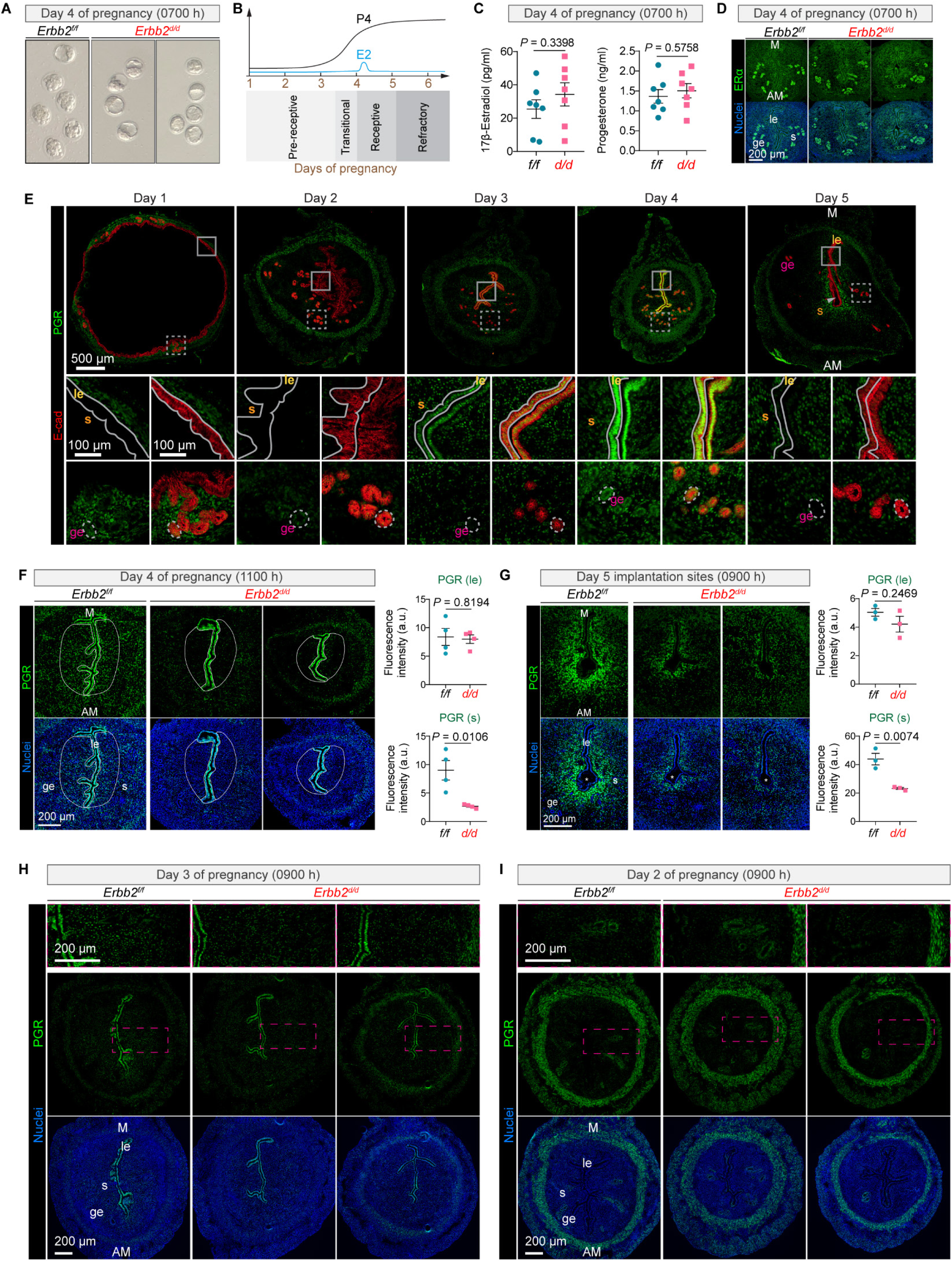
*Erbb2* deletion reduces stromal PGR expression. (**A**) Representative images of blastocysts flushed from the uteri of *Erbb2^f/f^*(n = 3) and *Erbb2^d/d^* (n = 3) mice at 0700 h on day 4 of pregnancy. **(B)** Schematic diagram illustrates the dynamic profiles of serum estradiol (E_2_) and progesterone (P_4_) levels during early pregnancy in mice. The shaded bars at the bottom indicate the corresponding phases of uterine sensitivity: pre-receptive, transitional, receptive, and refractory. (**C**) Serum levels of 17β-estradiol and progesterone measured at 1100 h on day 4 of pregnancy between *Erbb2^f/f^* (n = 7) and *Erbb2^d/d^* (n = 7) mice. Data are presented as mean ± SEM. (**D**) Immunofluorescence of ERα of *Erbb2^f/f^* (n = 6) and *Erbb2^d/d^*(n = 6) uteri at 1100 h on day 4 of pregnancy. Scale bar, 200 µm. (**E**) Representative images showing the expression pattern of PGR in mouse uteri on days 1 to 5 of pregnancy. n = 3 for each time point. Scale bars, 500 µm; magnified scale bars, 100 µm. le luminal epithelium, ge glandular epithelium, s stroma, M mesometrial pole, AM antimesometrial pole. (**F**) Immunofluorescence of PGR in uterine sections collected at 1100 h on day 4 of pregnancy from *Erbb2^f/f^*(n = 4) and *Erbb2^d/d^*(n = 4) mice. Scale bar, 200 µm. White outlines mark the uterine sub-luminal stromal area. (**G**) Immunofluorescence of PGR at 0900 h on day 5 of pregnancy in implantation sites from *Erbb2^f/f^* (n = 3) and *Erbb2^d/d^* (n = 3) mice. Scale bar, 200 µm. Asterisks indicate the location of the embryo. (**H**) Immunofluorescence of PGR at day 3 of pregnancy (0900 h) from *Erbb2^f/f^* (n = 3) and *Erbb2^d/d^* (n = 3) uteri. Scale bar, 200 µm; magnified scale bar: 200 µm. (**I**) Immunofluorescence of PGR at day 2 of pregnancy (0900 h) from *Erbb2^f/f^* (n = 3) and *Erbb2^d/d^* (n = 3) uteri. Scale bar, 200 µm; magnified scale bar: 200 µm. le luminal epithelium, ge glandular epithelium, s stroma, M mesometrial pole, AM antimesometrial pole.

PGR expression follows a spatiotemporal pattern across the peri-implantation uterus. Luminal epithelial PGR rises from day 3, peaks on day 4, and recedes by day 5; as it recedes, stromal PGR emerges on day 3 and persists through day 5, while glandular epithelial PGR remains detectable before implantation (Figure 6E), indicating that compartment-specific PGR dynamics coordinate the transition of the uterus from the receptive to the implantation phase.

From day 4, PGR is expressed in both the luminal epithelial and subluminal stromal cells of *Erbb2^f/f^* uteri, but stromal expression is reduced upon *Erbb2* loss while epithelial expression remains intact (Figure 6F). Following implantation on day 5, PGR expression expands from the primary decidual zone (PDZ) to the secondary decidual zone (SDZ) as decidualization progresses in *Erbb2^f/f^* uteri. In contrast, *Erbb2* deficiency reduces PGR expression on both day 5 and day 6 (Figure 6G; Supplemental Figure 1B). Having established that stromal PGR in *Erbb2^d/d^* uteri fails to reach *Erbb2^f/f^* levels, we next asked whether this difference is detectable at earlier stages. Because stromal PGR expression rises from day 3, we examined uteri on days 2 and 3, where only subtle differences distinguish *Erbb2^f/f^* and *Erbb2^d/d^* uteri (Figure 6H and I). A hallmark of day 2 is apoptosis in the uterine luminal epithelium, which resolves by day 3 (30). We next examined whether uterine deletion of *Erbb2* affects this process, as assessed by cleaved caspase-3 staining. In *Erbb2^f/f^* uteri, cleaved caspase-3 signals are evident in luminal epithelial cells on day 2 and decline by day 3. Apoptosis is comparable between *Erbb2^f/f^*and *Erbb2^d/d^* uteri on days 2 and 3 (Supplemental Figure 2A and B), indicating that the defect is specific to stromal PGR rather than early epithelial remodeling. Taken together, these findings demonstrate that *Erbb2* governs stromal PGR from the receptive phase through implantation and decidualization.

### Extended progesterone supplementation rescues implantation in *Erbb2^d/d^* mice

If reduced stromal PGR is what defers implantation in *Erbb2^d/d^* uteri, then raising circulating progesterone should restore it. We tested this directly, administering oil or progesterone from day 2 to day 4 of pregnancy and collecting uteri on day 6. Three days of progesterone failed to induce the clear blue bands seen in *Erbb2^f/f^* mice (Figure 7A), and stromal PGR in *Erbb2^d/d^* uteri remained low despite the supplementation. The decidual program was equally refractory: HAND2 and *Bmp2* expression in progesterone-treated *Erbb2^d/d^* uteri remained below *Erbb2^f/f^* levels and comparable to oil-treated *Erbb2^d/d^* uteri (Figure 7B). Thus, supplementing progesterone across the receptive window restores neither stromal PGR nor decidualization–the defect in *Erbb2^d/d^* uteri is not one of progesterone abundance.

**Figure 7.**
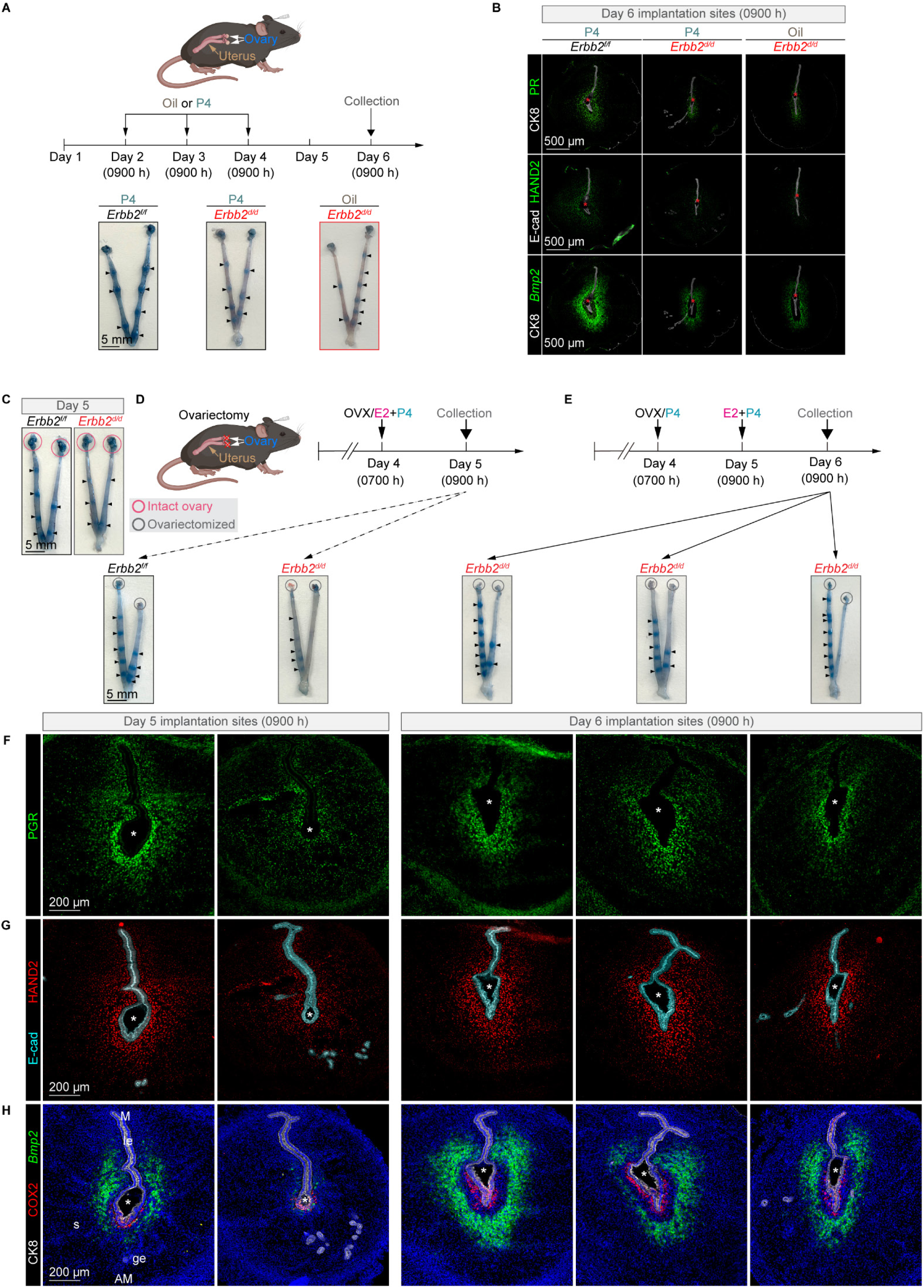
One-day P4 supplementation rescues deferred implantation in *Erbb2^d/d^* mice. (**A**) Schematic diagram of the three-day P_4_ rescue experiment. Mice (*Erbb2^f/f^* with P_4_ treatment: n = 3; *Erbb2^d/d^* with P_4_ treatment: n = 3; *Erbb2^d/d^* with oil treatment: n = 3) received subcutaneous injections of P_4_ or vehicle (oil) from day 2 to day 4 of pregnancy, and uteri were harvested on day 6. Scale bar, 5 mm. (**B**) Immunofluorescence of PGR and CK8, HAND2 and E-cad, and FISH of *Bmp2* and immunofluorescence of CK8 in *Erbb2^f/f^*and *Erbb2^d/d^* mice at day 6 implantation sites. (*Erbb2^f/f^* with P_4_ treatment: n = 3; *Erbb2^d/d^* with P_4_ treatment: n = 3; *Erbb2^d/d^* with oil treatment: n = 3). Scale bar, 500 μm. (**C**) Representative images of implantation sites in *Erbb2^f/f^* and *Erbb2^d/d^* mice at day 5 of pregnancy. (**D**) Schematic representation of the extended progesterone priming model under ovariectomized conditions. Ovariectomized (OVX) mice (n = 3 for *Erbb2^f/f^*, n = 3 for *Erbb2^d/d^*) were treated with E_2_ and P_4_ on day 4 (0700 h), followed by tissue collection at 0900 h on day 5. To determine whether extended progesterone priming could rescue the deferred implantation observed in *Erbb2^d/d^*mice, ovariectomized *Erbb2^d/d^* (n = 3) mice were first treated with P_4_ alone for 24 hours on day 4 (0700 h), then with combined E_2_ and P_4_ for another 24 hours on day 5 at 0900 h and collected on day 6 (0900 h). Blue bands indicate implantation sites. (**E**) Immunofluorescence PGR shows decreased expression in OVX *Erbb2^d/d^* uteri following immediate E_2_+P_4_ treatment, but enhanced expression after prior P4 exposure. Scale bar, 200 μm. (**F**) Immunofluorescence of HAND2 and E-cadherin (E-cad) shows increased HAND2 expression in stromal cells of *Erbb2^d/d^* uteri after P_4_ pre-treatment. Scale bar, 200 μm. (**G**) FISH of *Bmp2* and immunofluorescence of COX2 and CK8 across different groups. DAPI labels nuclei. Scale bar, 200 μm. Asterisks indicate embryo location. le luminal epithelium, ge glandular epithelium, s stroma, M mesometrial pole, AM antimesometrial pole.

If the amount of progesterone is not limiting, its timing may be. We therefore asked whether delivering P4 one day ahead of implantation, rather than across the receptive window, could rescue the defect (Figure 7C). To impose precise temporal control over hormone exposure, we turned to a delayed and activated implantation model. Ovariectomized *Erbb2^f/f^* and *Erbb2^d/d^* mice given E2 and P4 reproduced on-time implantation on day 5 of pregnancy (Figure 7D). When ovariectomized *Erbb2^d/d^* mice received P4 alone for 24 hours before combined E2 and P4 activation, visible blue bands returned (Figure 7E), and the molecular hallmarks of receptivity followed: PGR remained low immediately after E2 and P4 injection but rose when P4 preceded the combined treatment (Figure 7F), and HAND2 and *Bmp2* were induced upon P4 pre-treatment, signaling a restored decidual response (Figure 7G and H). COX2 remained comparable across groups, indicating that embryo attachment itself proceeded normally (Figure 7H). A single day of P4 priming ahead of E2 activation thus restores PGR signaling and reinstates on-time implantation in *Erbb2^d/d^* mice–placing the requirement for *Erbb2* not in the level of progesterone but in the timing of its priming.

### *Erbb2* is required for uterine gland LIF signaling and receptivity

Having traced the defect to the timing of progesterone priming, we next asked how *Erbb2* loss reshapes the receptive uterus itself. Two uterine receptivity markers, *Msx1* (estrogen and progesterone unresponsive) (31) and *Ihh* (progesterone responsive) (32, 33) were examined. On day 4, *Msx1* is enriched in the uterine glands over the luminal epithelium in *Erbb2^f/f^* uteri. However, *Erbb2^d/d^* uteri show comparable expression in both luminal and glandular epithelium (Supplemental Figure 3A). Additionally, FISH of *Ihh* reveals comparable expression levels in *Erbb2^f/f^* and *Erbb2^d/d^* uteri (Supplemental Figure 3B). On day 4 of pregnancy, luminal and glandular epithelial cells cease proliferation, while stromal cells begin to proliferate (34). In contrast, *Erbb2^d/d^* epithelial cells fail to exit the cell cycle and continue to proliferate, as marked by Ki67 (Supplemental Figure 3C).

Gland morphogenesis is an important event during uterine development (35). We first asked whether the glands form normally before pregnancy. At postnatal day 30, which approximates the onset of the estrous cycle in these *Erbb2* strains, uterine glands appear as simple elongated structures that are comparable between *Erbb2^f/f^*and *Erbb2^d/d^* mice, as shown by E-cadherin and FOXA2 co-staining (Supplemental Figure 4), indicating that early gland formation proceeds independently of *Erbb2*. During early pregnancy, they gradually undergo branching morphogenesis from day 1 to day 4 to establish uterine receptivity (36–38). The defect emerges during pregnancy: 3D imaging revealed that *Erbb2^d/d^* glands fail to undergo normal branching morphogenesis, appearing structurally disorganized relative to *Erbb2^f/f^*uteri (E-cadherin and FOXA2 co-staining; Supplemental Figure 5A). LIF is specifically expressed in uterine glands on day 4 and is critical to embryo implantation (39, 40). We found that *Lif* is primarily expressed in the uterine glands on day 4 of pregnancy; however, FISH and qPCR results show that its expression was reduced in *Erbb2^d/d^* uterine glands (Supplemental Figure 5B and C). We previously showed that the *Prss29+* glandular cells predominantly produce LIF on day 4 (36). *Prss29* mRNA, marking the glandular cells that are the predominant source of LIF, was likewise decreased in *Erbb2^d/d^*glands (FISH and qPCR; Supplemental Figure 5D and E), pointing to a loss of the LIF-producing glandular population.

To test whether reduced glandular LIF underlies the receptivity defect, we asked whether restoring LIF could rescue it. Recombinant LIF (rLIF) was injected intraperitoneally into *Erbb2^f/f^* and *Erbb2^d/d^*mice on day 4 of pregnancy (Supplemental Figure 6A). rLIF induced clear blue bands in both genotypes, confirming that embryo attachment took place; the implantation sites in *Erbb2^d/d^*mice, however, remained smaller than in *Erbb2^f/f^* mice (Supplemental Figure 6B), indicating that attachment occurred late. This delay carried forward: rLIF did not restore PGR expression in *Erbb2^d/d^* uteri (Supplemental Figure 6C), and the marginal rise in the decidualization marker *Bmp2* fell short of a normal decidual response (Supplemental Figure 6D). Restoring LIF therefore cannot rescue implantation or decidualization in *Erbb2^d/d^* mice, locating the essential requirement in stromal PGR rather than glandular LIF.

### Endometrial PGR was decreased in patients with recurrent spontaneous abortion

Our mouse data trace deferred implantation to a failure of stromal progesterone priming; we next asked whether the same stromal deficit marks human pregnancy loss. Mining a single-cell transcriptomic dataset of endometrium from women with recurrent spontaneous abortion (RSA) and fertile controls, we found PGR selectively downregulated in the stromal compartment (Figure 8A and B). The reduction was concentrated in the decidual stromal (DS) clusters, whereas PGR in epithelial and other compartments was largely unaffected.

**Figure 8.**
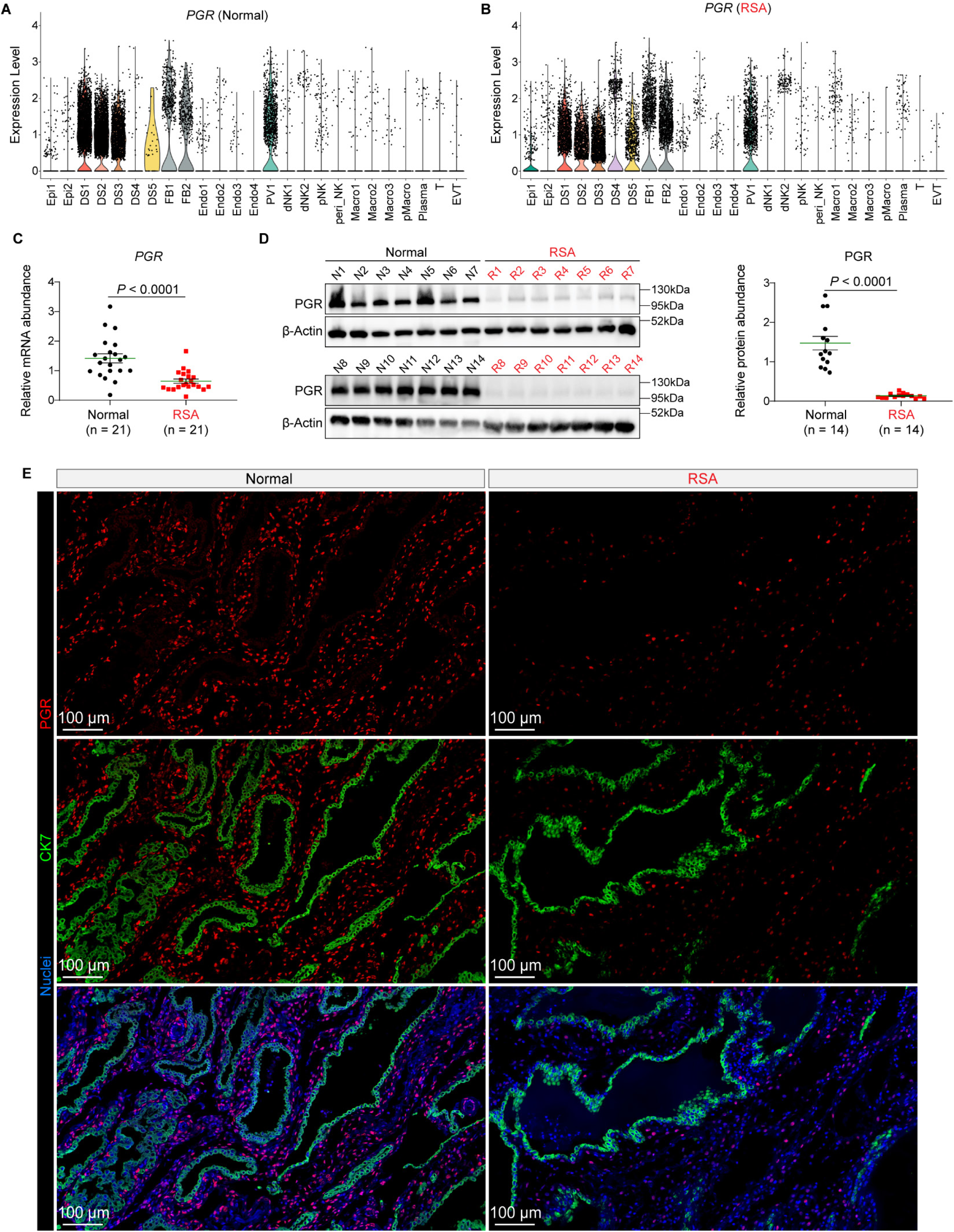
Endometrial stromal PGR expression is downregulated in patients with recurrent spontaneous abortion. (**A** and **B**) Violin plots showing the expression levels of PGR across various cell populations identified by single-cell RNA sequencing Normal (**A**) and RSA patients (**B**). (**C**) Quantitative RT-PCR validation of PGR mRNA abundance in endometrial tissues from an independent cohort of Normal (n = 21) and RSA (n = 21) patients. (**D**) Representative Western blot and densitometric quantification of PGR protein levels in Normal and RSA patients. β-actin served as the loading control. (**E**) Representative immunofluorescence staining of PGR (red) and the epithelial marker cytokeratin 7 (CK7, green) in Normal and RSA patients. Nuclei were counterstained with DAPI (blue). Scale bars: 100 μm. Data in (**C**) and (**D**) are presented as mean ± SEM. Each dot represents an individual sample. *P* values were determined by unpaired 2-tailed Student’s *t* test.

To validate this finding, we analyzed endometrial biopsies from an independent clinical cohort of RSA patients (n = 21) and fertile controls (n = 21). Consistent with the transcriptomic data, qPCR analysis confirmed a significant reduction in PGR mRNA abundance in RSA samples (Figure 8C). This transcriptional deficit translated to the protein level, as Western blot analysis demonstrated markedly lower PGR protein expression in RSA endometria (Figure 8D). Immunofluorescence indicated that this loss was largely stroma-restricted: epithelial PGR appeared unchanged, while nuclear PGR in the stromal compartment was reduced in RSA patients (Figure 8E). This pattern is consistent with the stroma-biased reduction of PGR observed in *Erbb2*-deficient mice. Together, the mouse and human data converge on stromal progesterone responsiveness as a shared determinant of implantation success and point to the ERBB2-PGR axis as a candidate contributor to recurrent pregnancy loss.

## Discussion

Embryo implantation requires communication between a competent blastocyst and the receptive uterus. In this study, we identify *Erbb2* as a critical regulator of stromal progesterone responsiveness, ensuring on-time implantation and appropriate decidualization. Our findings emphasize that stromal preparation is an essential determinant of uterine receptivity. Although *Erbb2* deletion did not affect ovarian steroid hormone levels, embryo development, or epithelial PGR expression, a marked reduction in stromal PGR emerged at the onset of the receptive window on day 4 and persisted at least through day 6. This defect triggered a series of abnormalities, including deferred implantation, impaired gland-crypt assembly, attenuated decidualization, and increased embryo resorption. The observation that epithelial-specific deletion of *Erbb2* resulted in normal fertility further supports that *Erbb2* functions predominantly in the stromal compartment during early pregnancy.

A central conclusion of this work is that *Erbb2* is necessary to establish a progesterone-responsive stromal environment. While epithelial PGR expression remained intact, stromal PGR failed to rise appropriately and remained diminished throughout implantation and decidual progression. Because stromal PGR governs the induction of key mediators such as HAND2, which suppresses epithelial proliferation and promotes stromal differentiation, reduced PGR in *Erbb2^d/d^* uteri readily explains both the heightened epithelial proliferation and the loss of HAND2 around the implanting embryo. These findings place *Erbb2* upstream of stromal PGR and highlight its role in coordinating epithelial-stromal communication.

Deferred implantation has previously been observed in *Pla2g4a* (41) and *Hbegf* (25) mutant mice. Similarly, our study demonstrates that *Erbb2*-deficient uteri undergo deferred, rather than failed, implantation. The absence of blue dye reactions on day 4.5 and the presence of unattached blastocysts within the uterus indicate a delay in the attachment reaction. Consistent with this phenotype, both *Hbegf* and COX2 were diminished in *Erbb2^d/d^* uteri at day 4.5, supporting the conclusion that defective stromal preparation contributes to delayed embryo-uterine communication. Notably, progesterone administration one day earlier in a delayed implantation model restored stromal PGR, HAND2, *Bmp2*, and implantation timing. This experiment demonstrates that the temporal window of progesterone administration is a critical determinant of stromal competency, and further positions *Erbb2* as a regulator of this temporal sensitivity.

Beyond its stromal functions, *Erbb2* also influences uterine gland architecture and activity. Abnormal gland morphology, decreased Lif and Prss29 expression, and disrupted gland-crypt assembly were all evident in *Erbb2^d/d^* uteri. Although exogenous LIF partially enhanced *Bmp2* levels, it did not restore stromal PGR or fully rescue implantation and decidualization, indicating that *Erbb2* contributes to uterine receptivity primarily through a PGR-dependent stromal pathway, rather than through gland-derived LIF alone. The 3D imaging data further reveal that *Erbb2* integrates epithelial, stromal, and glandular interactions required for proper embryo implantation.

A stroma left poorly prepared by reduced PGR should carry a molecular signature well before the preimplantation estrogen surge. Profiling the day-4 endometrial transcriptome revealed such a signature: while classical receptivity markers such as *Msx1* and *Ihh* were unchanged, the TGF-β superfamily member Growth differentiation factor 7 (*Gdf7*) was selectively reduced in *Erbb2^d/d^* uteri (Supplemental Figure 7). Previous studies have identified *Gdf7* as an oviduct- and uterus-enriched mesenchymal secreted factor (42, 43), and stromal GDF7 was decreased in human endometrioid endometrial cancer by scRNA-seq data (44, 45). *Gdf7* localizes to the subluminal stroma on day 4 and is diminished there in *Erbb2^d/d^*uteri (Supplemental Figure 8), so its loss tracks the localized failure of stromal PGR at the very compartment and stage where receptivity is established. *More broadly*, *Gdf7* (42) dysregulation has been linked to human endometrial pathologies, including intrauterine adhesions (46) and endometriosis (43). Whether *Gdf7* actively contributes to implantation awaits lineage-specific deletion; for now, its stromal-specific downregulation marks *Gdf7* as a candidate readout of stromal preparation downstream of the ERBB2-PGR axis.

The relevance of our findings to recurrent spontaneous abortion (RSA) is supported by reanalysis of published single-cell RNA-seq datasets demonstrating reduced *ERBB2* and PGR expressions in decidual stromal clusters from RSA patients. These observations were validated by qPCR, western blotting, and immunofluorescence. Although these data do not establish causality, they suggest that altered ERBB2-PGR axis may contribute to suboptimal stromal function in recurrent spontaneous abortion. Together, these results support a model in which ERBB2 preserves stromal progesterone responsiveness, enabling timely activation of HAND2, *Bmp2*, and stromal-gland communication required for embryo attachment and decidualization. Loss of *Erbb2* impairs these coordinated events, leading to deferred implantation, impaired decidualization, and adverse pregnancy outcomes.

Despite these advances, several questions remain. The transcriptional and epigenetic mechanisms by which *ERBB2* regulates stromal PGR expression remain undefined. Whether *ERBB2* influences PGR transcription, protein stability, or cofactor recruitment requires further investigation. In addition, functional studies using *Gdf7* gain- and loss-of-function models will be important to determine whether *Gdf7* marks stromal readiness or actively contributes to implantation. Finally, the absence of stromal-specific *Cre* drivers limits our ability to dissect stromal-autonomous functions of ERBB2. Developing more refined genetic tools will be essential for resolving the precise cell type-specific roles of ERBB2 during early pregnancy.

## Methods

### Mice

*Erbb2^f/f^* (47) mice were crossed with *Pgr^Cre/+^* (48) or *Ltf^Cre/+^* (49) mice to generate *Erbb2^f/f^ Pgr^Cre/+^* (*Erbb2^d/d^*) and *Erbb2^f/f^ Ltf^Cre/+^* (*Erbb2^epi/epi^*) mouse lines. These three genotypes of mice were housed in the animal care facility at Cincinnati Children’s Hospital Medical Center according to the National Institute of Health and institutional guidelines for laboratory animals. All protocols were approved by the Cincinnati Children’s Animal Care and Use Committee. Mice were provided with irradiated Laboratory Rodent Diet 5R53 and autoclaved water ad libitum. These mice were housed under a 12:12 hr light:dark cycle. At least three mice from each genotype were used for each individual experiment.

### Analysis of pregnancy events

Adult females from each genotype were randomly chosen and housed with a fertile WT male of choice overnight; the morning of finding a vaginal plug was considered successful mating (day 1 of pregnancy). To confirm that plug-positive mice were pregnant on day 4 of pregnancy, one uterine horn was flushed with saline to detect blastocyst existence. For day 4.5, day 5 and day 6 pregnant mice, 100 μl of 1% Chicago blue in saline were injected intravenously for 4 minutes to visualize implantation sites as blue bands, whether clear or faint. If no blue band was observed, uterine horns were flushed to check for the presence of embryos.

### Human sample collection

The tissue samples in this study were collected after obtaining written informed consent from all participants, with approval from the Institutional Ethics Committee of Chongqing Medical University. Participants in the recurrent spontaneous abortion (RSA) group met stringent inclusion criteria: 1) ≥2 episodes of unexplained spontaneous abortions; 2) confirmed normal karyotypes in both parents and abortus; and 3) exclusion of uterine anomalies, endocrine/metabolic disorders, autoimmune diseases, or infections (14 for western blot, 21 for qPCR). Healthy controls consisted of women undergoing elective termination of uncomplicated pregnancies without prior miscarriage history (14 for western blot, 21 for qPCR). RSA specimens were collected through ultrasound-guided curettage immediately after missing abortion diagnosis. Decidual tissues were macroscopically identified and processed as follows: samples were thoroughly washed with ice-cold PBS to remove blood residues, then flash-frozen in liquid nitrogen for subsequent analyses. Control specimens followed identical collection and processing protocols. All procedures were performed under strict sterile conditions using pre-chilled reagents to maintain sample integrity. Demographic characteristics of the cohorts are presented in Supplemental Table 1.

### Western Blot

Total protein was isolated using lysis buffer, and concentrations were determined with a BCA protein assay kit (Thermo Pierce Biotechnology, USA) following the manufacturer’s protocol. Equal amounts of protein were resolved by SDS-PAGE at 100 V for 120 min and subsequently electrotransferred to PVDF membranes (0.22 μm pore size; Millipore, Germany) at 250mA for 120 min. The membranes were blocked with 5% skim milk (Bio-Rad, USA) and incubated overnight at 4 °C with the indicated primary antibodies: anti-ERBB2 (2165S, 1:1000, Cell Signaling Technology), anti-PGR (ab32085,1:1000, Abcam) and anti-β-Actin (HRP-66009, 1:1000, Proteintech). After washing, membranes were incubated for 1 h at room temperature with horseradish peroxidase (HRP)-linked anti-rabbit secondary antibodies (1:5000, Santa Cruz Biotechnology). Immunoreactive bands were visualized using an enhanced chemiluminescence reagent (Advansta, USA) and imaged with a Bio-Rad XRS+ system, and densitometric analysis was performed using Bio-Rad Image Lab software (version 5.2). Primary antibodies were diluted in 5% BSA prepared in 1× TBS containing 0.1% Tween-20.

### Quantitative PCR (qPCR)

Total RNA was isolated using TRIzol reagent (Invitrogen, USA). First-strand cDNA was then generated with the Transcriptor First Strand cDNA Synthesis Kit (Roche, Germany) following the supplier’s protocol. Quantitative PCR was carried out using FastStart Essential DNA Green Master Mix (Roche, Germany) on a Bio-Rad CFX96 real-time PCR platform. The qPCR program consisted of an initial step at 95 °C for 10 min to activate the polymerase and denature the template, followed by 40 amplification cycles of 95 °C for 10 s, annealing for 30 s at the primer-specific optimal temperature, and extension at 72 °C for 10 s. Melt-curve analysis was subsequently performed by increasing the temperature from 65 °C to 95 °C in 0.5 °C steps with a 5 s hold at each increment. Data processing was conducted using Bio-Rad CFX Maestro software (version 1.1). Target transcript levels were quantified based on Ct values, and relative expression was calculated after normalization to the internal reference gene β-actin. Primer sequences for all genes analyzed are provided in Supplemental Table 2.

### Histology

Tissue sections from control and experimental groups were processed on the same slides. Frozen sections (12 μm) were fixed in 4% PFA for 10 min at room temperature and then stained with hematoxylin and eosin for light microscopy analysis.

### Fluorescence *In Situ* Hybridization (FISH)

FISH experiments were performed as previously described (40). Briefly, frozen sections (12 µm) were processed on the same slide for each probe. Following fixation (in 4% paraformaldehyde) and acetylation, slides were hybridized at 55°C with digoxigenin-labeled *Lif*, *Prss29*, *Msx1*, *Ihh*, *Hbegf*, *Ptgs2* or *Bmp2* probes. Anti-DIG-peroxidase was applied onto hybridized slides following washing and peroxide quenching. The color was developed by TSA (Tyramide Signal Amplification) fluorescein according to the manufacturer’s instructions (PerkinElmer). Epithelia were stained using a CK8 antibody (TROMA-I, DSHB). Glands were stained using a FOXA2 antibody (8186s, Cell Signaling Technology). Images were captured using a confocal microscope (Nikon Eclipse TE2000). Nuclei staining was performed using Hoechst 33342 (2 µg/ml, H1399, Thermo Scientific).

### Immunofluorescence (IF)

Frozen sections (12 µm) from each genotype were mounted and processed onto same slides. IF for anti-ERα (sc-542, 1:300, Santa Cruz), anti-PGR (8757, 1:300, Cell signaling Technology), anti-HAND2 (AF3876, 1:300, R&D Systems) and anti-Ki67 (RM-9106-S, 1:300, Invitrogen) were performed using secondary antibodies conjugated with Alexa-conjugated 488, 594 or 647 antibodies (Jackson ImmunoResearch, 1:300). Nuclei staining was performed using Hoechst 33342 (2 µg/ml, H1399, Invitrogen). Images were captured using a confocal microscope (Nikon Eclipse TE2000).

### Measurement of serum E_2_ and P_4_ levels

Sera were collected on day 4 of pregnancy (1100 h), and hormone levels were measured by enzyme immunoassay kits (Estradiol ELISA Kit, 501890, Cayman) and (Progesterone ELISA Kit, 582601, Cayman) as previously described (31).

### Whole-mount immunostaining for 3D imaging

The whole-mount immunostaining with 3DISCO tissue clearing was performed as previously described (21). Briefly, uterine samples were fixed in Dent’s Fixative (Methanol:DMSO (4:1)) overnight in -20 °C and then washed with 100% methanol for three times. The samples were bleached with 3% H_2_O_2_ in methanol at 4 °C overnight to remove pigmentation. After washing in 1% PBS-T for 6 times with 1 hour each, samples were incubated with E-cadherin (3195s, 1:100, Cell Signaling Technology), E-cadherin (13-1900, 1:100, Invitrogen), FOXA2 (WRAB-1200, 1:100, Seven Hills), COX2 (12282s, 1:100, Cell Signaling Technology) or PECAM1 (AF3628, 1:100, R&D Systems) at 4 °C on a rotor for 7 days. After incubation, the samples were washed with 1% PBS-T six times for 1 hour each and incubated with Alexa-conjugated 488, 594 or 647 antibodies (1:200, Jackson Immuno Research) in a light-proof box for 4 days at 4 °C. After six washes in 1% PBS-T at room temperature, samples were dehydrated in 100% methanol for 1 hour and then cleared in benzyl alcohol/benzyl benzoate (BABB) solution for at least 1 hour in a light-proof box.

### 3D imaging and processing

3D pictures were acquired using a Nikon FN1 Upright Microscope. Samples were laid on slides, covered with BABB, and enclosed by cover slips for confocal imaging using a 10X objective with 8 µm Z-stack. All files were generated by Nikon elements and were imported into Imaris (version 10.1, Bitplane) for visualization and 3D reconstruction. To obtain the 3D structure of the tissue, the surface tool was utilized. To isolate a specific region of the tissue, the surface tool was manually used to segment the images, and the mask option was selected for subsequent pseudo-coloring. 3D images were generated using the snapshot tool.

### Primary stromal cells isolation and treatment

Stromal cells from day 4 of the pregnant uterus were collected by enzymatic digestion as described previously (30). Uteri from *Erbb2^f/f^* and *Erbb2^d/d^* mice on day 4 of pregnancy were split longitudinally and cut into small fragments. The tissue pieces were incubated with pancreatin (25 mg/mL, Sigma) and dispase (6 mg/ mL, Gibco) for 1h at 4°C, followed by 20 min at room temperature and 5 min at 37 °C to remove luminal epithelial sheets through washing. The remaining tissue fragments were digested with type IV collagenase (300 U/mL, Washington) for 30 min at 37 °C to release stromal cells. The resulting cell suspensions were filtered through a 70-μm nylon mesh to remove glands and epithelial cell clumps. Stromal cells were suspended in DMEM/F12 (Gibco) supplemented with 10% heat-inactivated FBS (Gibco), 50 units/mL penicillin, 50 μg/mL streptomycin, and 1.25 μg/mL fungizone (Pen Strep; Gibco). Cells were seeded into 12-well plates and cultured overnight before initiating in vitro decidualization. For in vitro decidualization, stromal cells were treated with estradiol-17β (E2, 10 nM) and progesterone (P4, 1 μM) in DMEM/F12 media supplemented with 2% charcoal-stripped FBS (vol/vol) for 3 and 6 days. Cells were harvested at designated time points for RNA extraction.

### LIF rescue experiment

The LIF rescue experiment was as previously described (42). Briefly, *Erbb2^f/f^* and *Erbb2^d/d^*mice on day 4 of pregnancy (0900 h) received a single intraperitoneal injection of recombinant LIF (20 μg/mouse in 100 μl saline). Two days after the LIF injection, implantation sites were examined by intravenous injection of a blue dye solution.

### P_4_ rescue experiment

To determine whether an additional day of progesterone priming could rescue implantation, pregnant *Erbb2^f/f^* and *Erbb2^d/d^*mice were ovariectomized on day 4 of pregnancy (0700 h). Two groups of *Erbb2^f/f^*and *Erbb2^d/d^* mice immediately received subcutaneous injection estradiol-17β (E2, 25 ng/100 ul) and progesterone (P4, 1 mg/100 μl). These mice were sacrificed and collected on day 5 of pregnancy (0900 h). A third group of *Erbb2^d/d^* mice received an initial subcutaneous injection of P4 (1 mg/100 μl) immediately after ovariectomy and were maintained for 24 hours. On day 5 of pregnancy (0900 h), these mice were injected with E2 (25 ng/100 μl) and P4 (1 mg/100 μl) and subsequently sacrificed on day 6 of pregnancy (0900 h). Implantation sites were visualized by intravenous injection of a blue dye solution.

To assess the impact of prolonged progesterone on embryo implantation, pregnant *Erbb2^f/f^* and *Erbb2^d/d^* mice received subcutaneous injections of progesterone (1 mg/mouse) for three consecutive days, beginning on day 2 of pregnancy. Mice were sacrificed on day 6 (0900 h), and implantation sites were visualized via intravenous injection of a blue dye solution.

### Bulk RNA-sequencing analysis

Total RNA was isolated from day 4 mouse endometria and used for paired-end RNA sequencing. Sequencing reads were aligned to the mm10 reference genome using HISAT2 with default parameters. Resulting SAM files were converted to coordinate-sorted BAMs using samtools. To retain uniquely mapped primary reads, alignments with *NH:i:1* and without secondary or supplementary flags were selected. Gene-level counts were generated with featureCounts (paired-end mode; chimeric and discordant pairs excluded) using an mm10-compatible GTF annotation. Differential expression analysis was performed on raw counts using DESeq2, with multiple-testing correction by the Benjamini-Hochberg method. Genes with adjusted *p* < 0.05 and, where applicable, |log2(fold change)| > 1 were considered differentially expressed. Variance-stabilized counts were used for principal component analysis and sample-sample correlation.

### Statistical analysis

Statistical analyses were conducted using GraphPad Prism (v8.0) and R (v4.4.1) along with RStudio (2024.04.2). Each experiment was repeated at least three times. Data are shown as mean ± SEM. Statistical analyses were performed using a two-tailed Student’s *t*-test. A *P* value less than 0.05 was considered statistically significant.

## Data availability

The sequencing data was submitted to National Center for Biotechnology Information (NCBI) BioProject database with accession number of PRJNA1364307. The raw and processed bulk RNA-seq data generated in this study have been deposited in the NCBI Gene Expression Omnibus (GEO) under accession number GSE310168.

## Author contributions

XFS, SKD, and HBQ designed research; BL, CZ, AD, XLL, and WBD performed research; BL, and XFS analyzed data; and BL and XFS wrote the paper.

## Acknowledgments

We are grateful to Francesco DeMayo (National Institute of Environmental Health Science, Research Triangle Park, NC) and John B. Lydon (Baylor College of Medicine, Houston, TX) originally provided *Pgr^Cre/+^* mice. We thank BioRender.com for providing the platform.

---

**Supplemental Figure 1.**
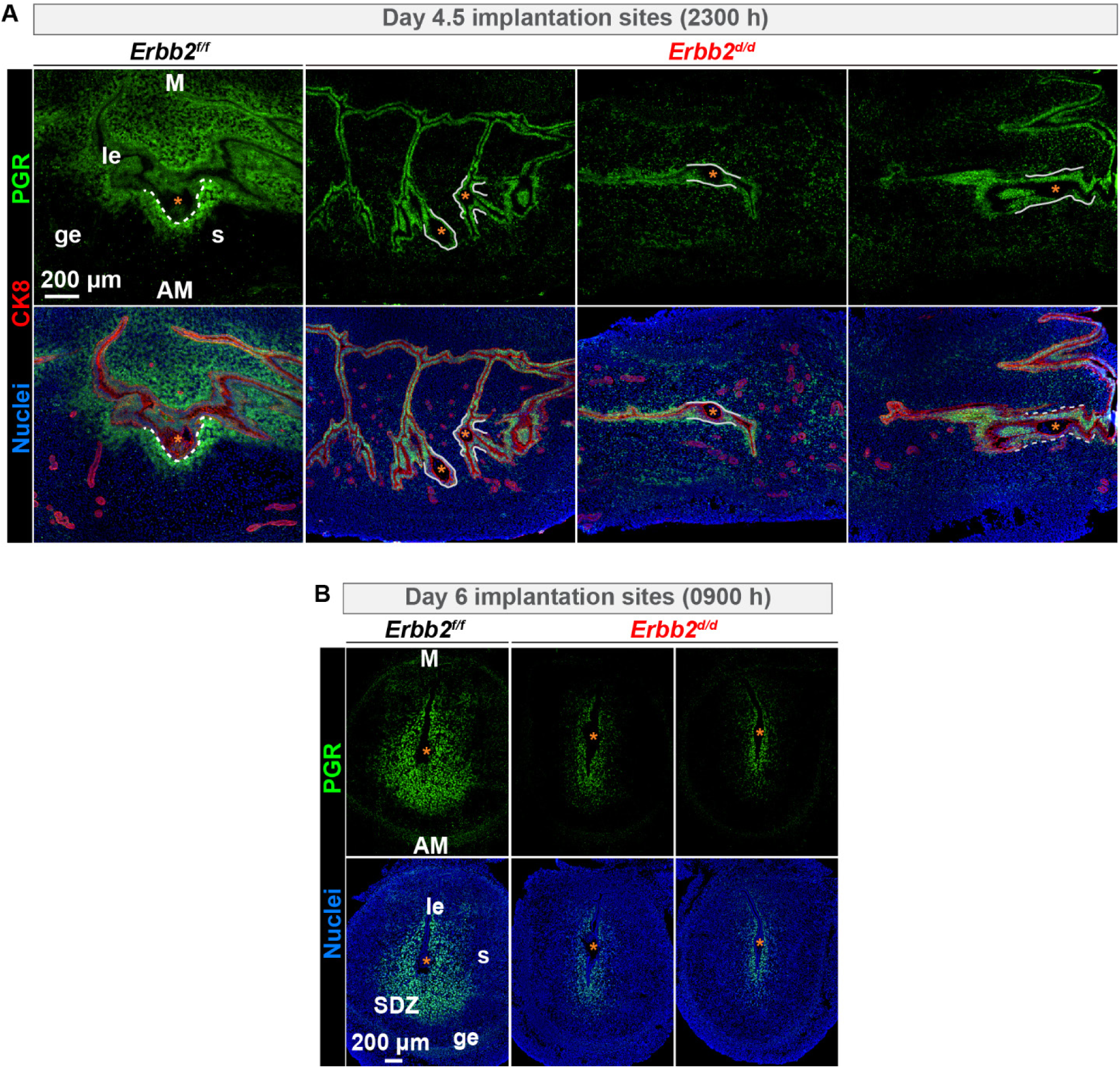
PGR expression in *Erbb2^f/f^* and *Erbb2^d/d^* uteri on day 2, day 3, day 4.5, and day 6 of pregnancy. (**A**) Immunofluorescence of PGR and CK8 in implantation sites at day 4.5 of pregnancy (2300 h) from *Erbb2^f/f^* (n = 3) and *Erbb2^d/d^* (n = 3) uteri. Scale bar, 200 µm. (**B**) Immunofluorescence of PGR in implantation sites at 0900 h on day 6 from *Erbb2^f/f^* (n = 4) and *Erbb2^d/d^* (n = 4) uteri. Scale bar, 200 µm. Asterisks indicate the location of the embryo. le luminal epithelium, ge glandular epithelium, s stroma, M mesometrial pole, AM antimesometrial pole, SDZ secondary decidual zone.

**Supplemental Figure 2.**
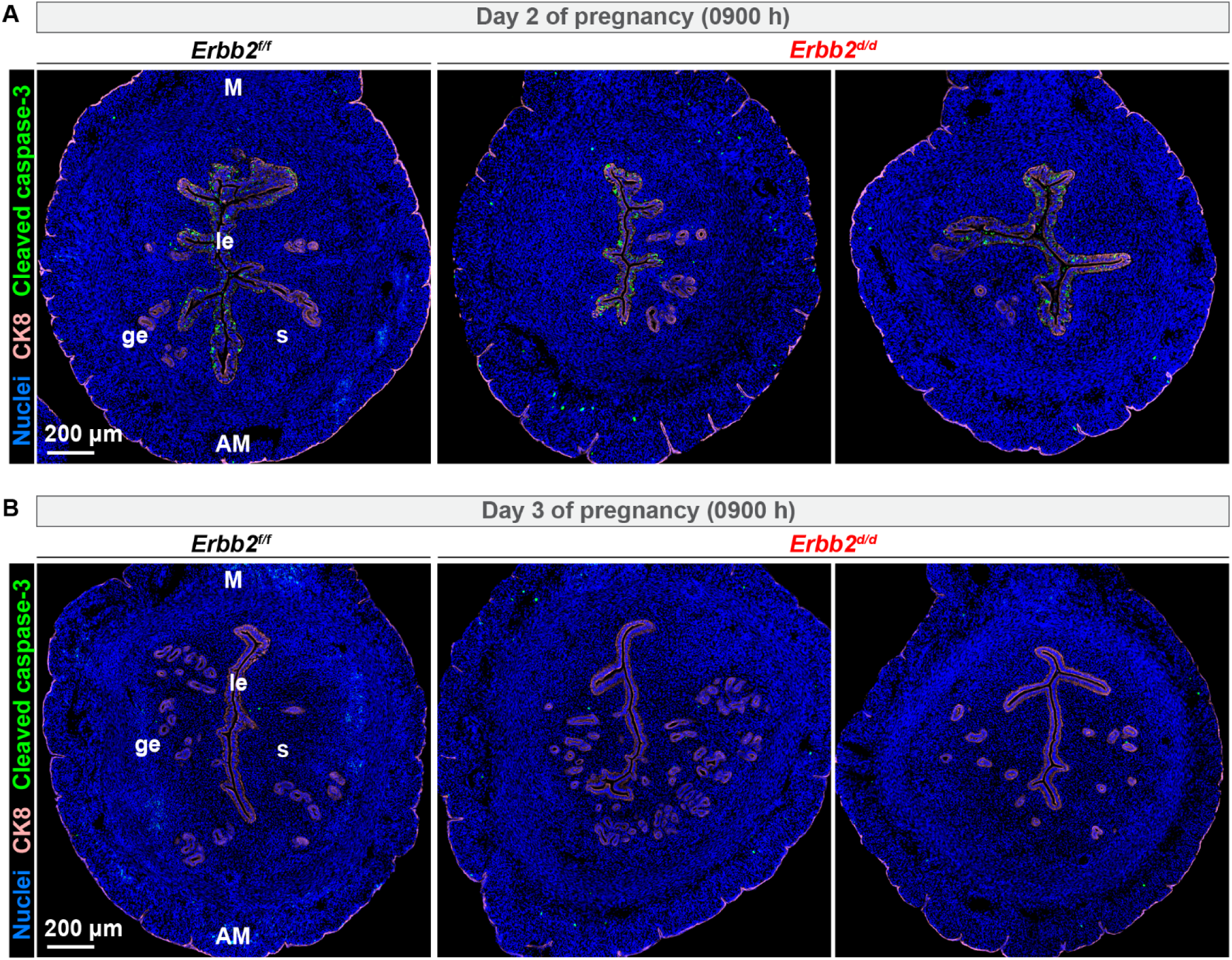
Analysis of epithelial apoptosis in *Erbb2^d/d^* mice on day 2 and day 3 of pregnancy. (**A** and **B**) Immunofluorescence staining of cleaved caspase-3 and CK8 from *Erbb2^f/f^* and *Erbb2^d/d^* mice on day 2 (**A**; n = 3 for *Erbb2^f/f^*, n = 3 for *Erbb2^d/d^*) and day 3 (**B**; n = 3 for *Erbb2^f/f^*, n = 3 for *Erbb2^d/d^*) of pregnancy (0900 h). Nuclei were counterstained with DAPI. le luminal epithelium, ge glandular epithelium, s stroma, M mesometrial pole, AM antimesometrial pole.

**Supplemental Figure 3.**
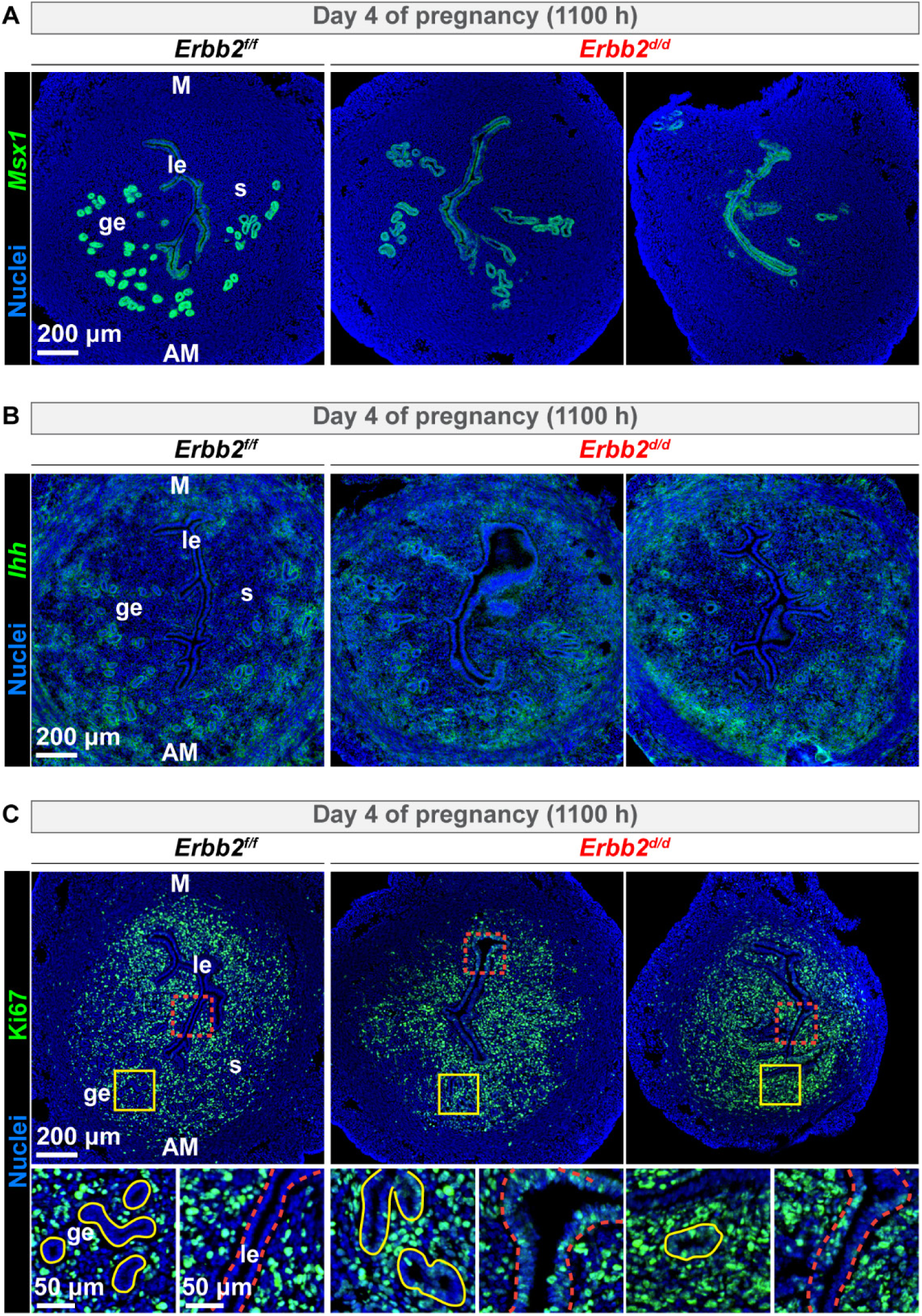
M*s*x1 and *Ihh* are not altered in *Erbb2^d/d^*uteri on day 4 of pregnancy. (**A**) FISH of *Msx1* from *Erbb2^f/f^* (n = 3) and *Erbb2^d/d^* (n = 4) mice on day 4 of pregnancy (1100 h). Scale bar, 200 μm. (**B**) FISH of *Ihh* revealed comparable expression levels in *Erbb2^f/f^* (n = 6) and *Erbb2^d/d^* (n = 6) uteri. Scale bar, 200 μm. (**C**) Ki67 immunostaining from *Erbb2^f/f^* (n = 3) and *Erbb2^d/d^*(n = 4) mice on day 4 of pregnancy (1100 h). Scale bar, 200 μm; magnified scale bars: 50 μm. le luminal epithelium, ge glandular epithelium, s stroma, M mesometrial pole, AM antimesometrial pole.

**Supplemental Figure 4.**
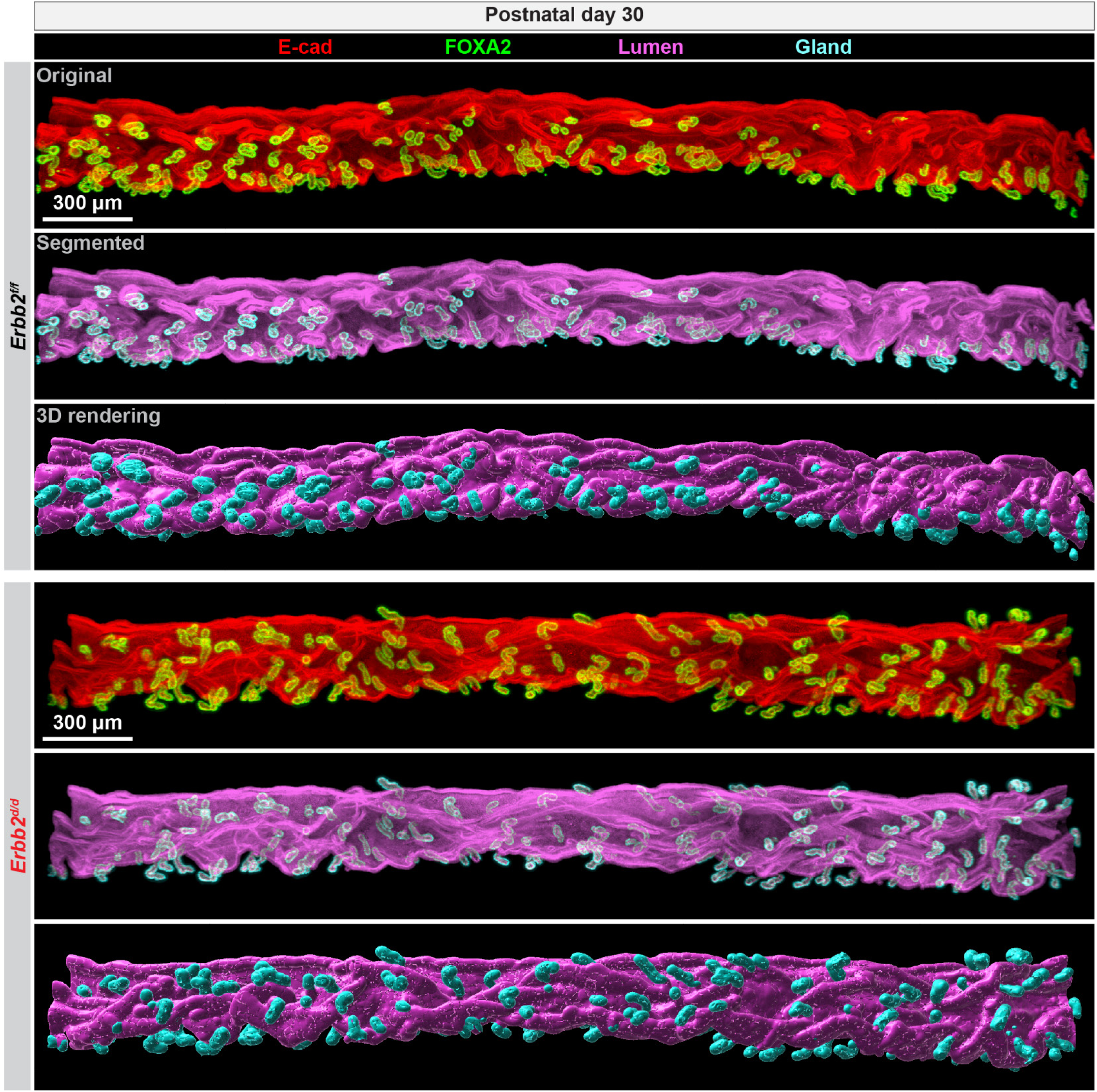
3D imaging from *Erbb2^f/f^* and *Erbb2^d/d^* females on postnatal day 30. Original double immunostaining with E-cadherin and FOXA2 and 3D rendering pictures. E-cadherin marks both luminal epithelial and glandular epithelial cells, while FOXA2 only marks glands. Scale bars, 300 μm.

**Supplemental Figure 5.**
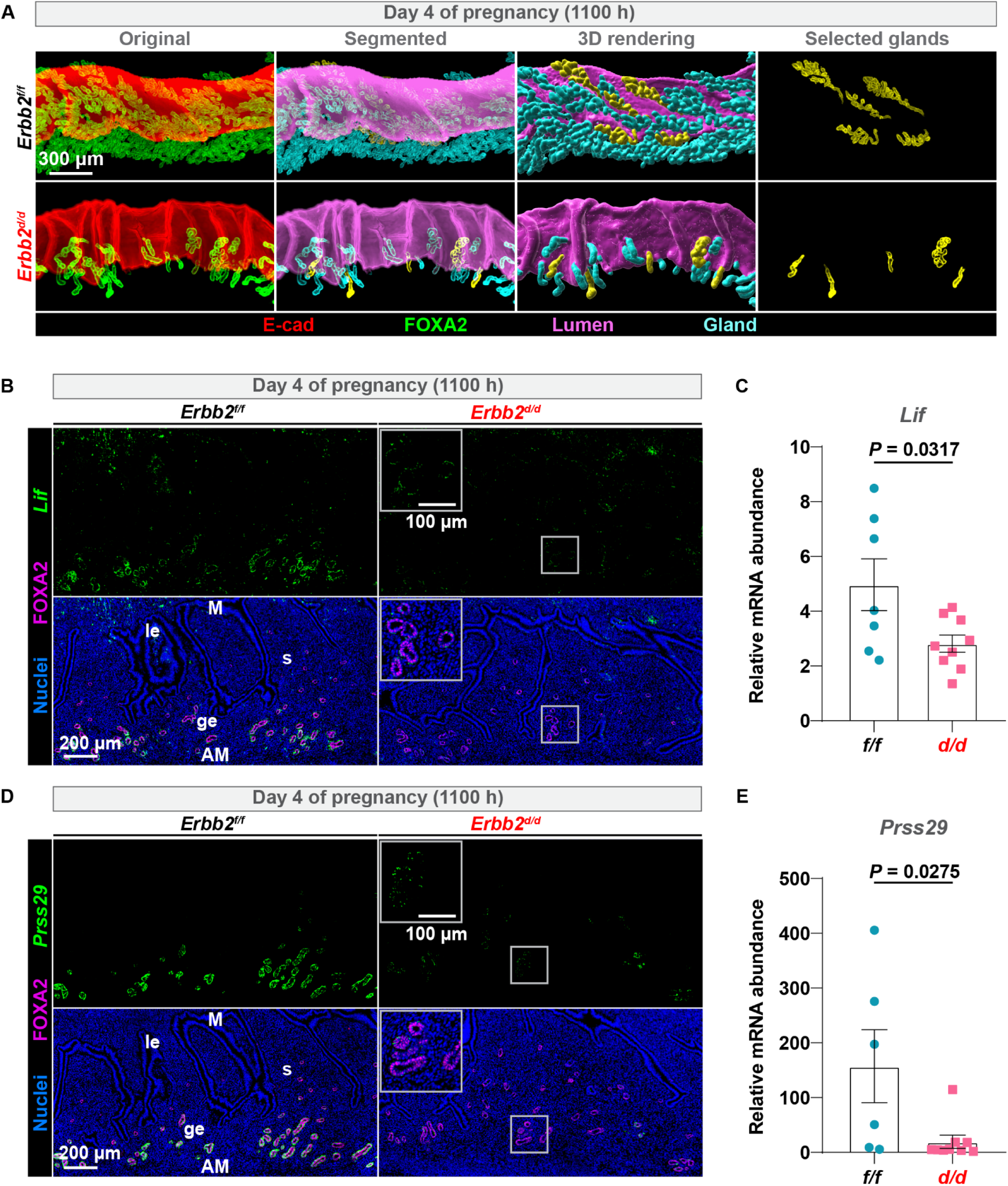
Uterine gland morphology and LIF secretion are abnormal in *Erbb2^d/d^* mice on day 4 of pregnancy. (**A**) 3D imaging in *Erbb2^f/f^* and *Erbb2^d/d^* uteri on day 4 of pregnancy at 1100 h. Scale bar, 300 μm. Original (E-cadherin and FOXA2), Segmented, and 3D rendering. le luminal epithelium, ge glandular epithelium, s stroma, M mesometrial pole, AM antimesometrial pole. (**B**) FISH of *Lif* and Immunofluorescence of FOXA2 in *Erbb2^f/f^* (n = 6) and *Erbb2^d/d^* (n = 7) uteri on day 4 of pregnancy at 1100 h. Scale bar, 200 μm; magnified scale bar: 100 μm. (**C**) qPCR analysis of *Lif* mRNA levels in uteri from *Erbb2^f/f^* (n = 7) and *Erbb2^d/d^* (n = 9) mice. Data are presented as mean ± SEM. (**D**) FISH of *Prss29* and Immunofluorescence of FOXA2 in *Erbb2^f/f^*(n = 6) and *Erbb2^d/d^* (n = 7) uteri on day 4 of pregnancy at 1100 h. Scale bar, 200 μm; magnified scale bar: 100 μm. (**E**) qPCR analysis of *Prss29* mRNA levels in uteri from *Erbb2^f/f^* (n = 6) and *Erbb2^d/d^* (n = 9) mice. Data are presented as mean ± SEM.

**Supplemental Figure 6.**
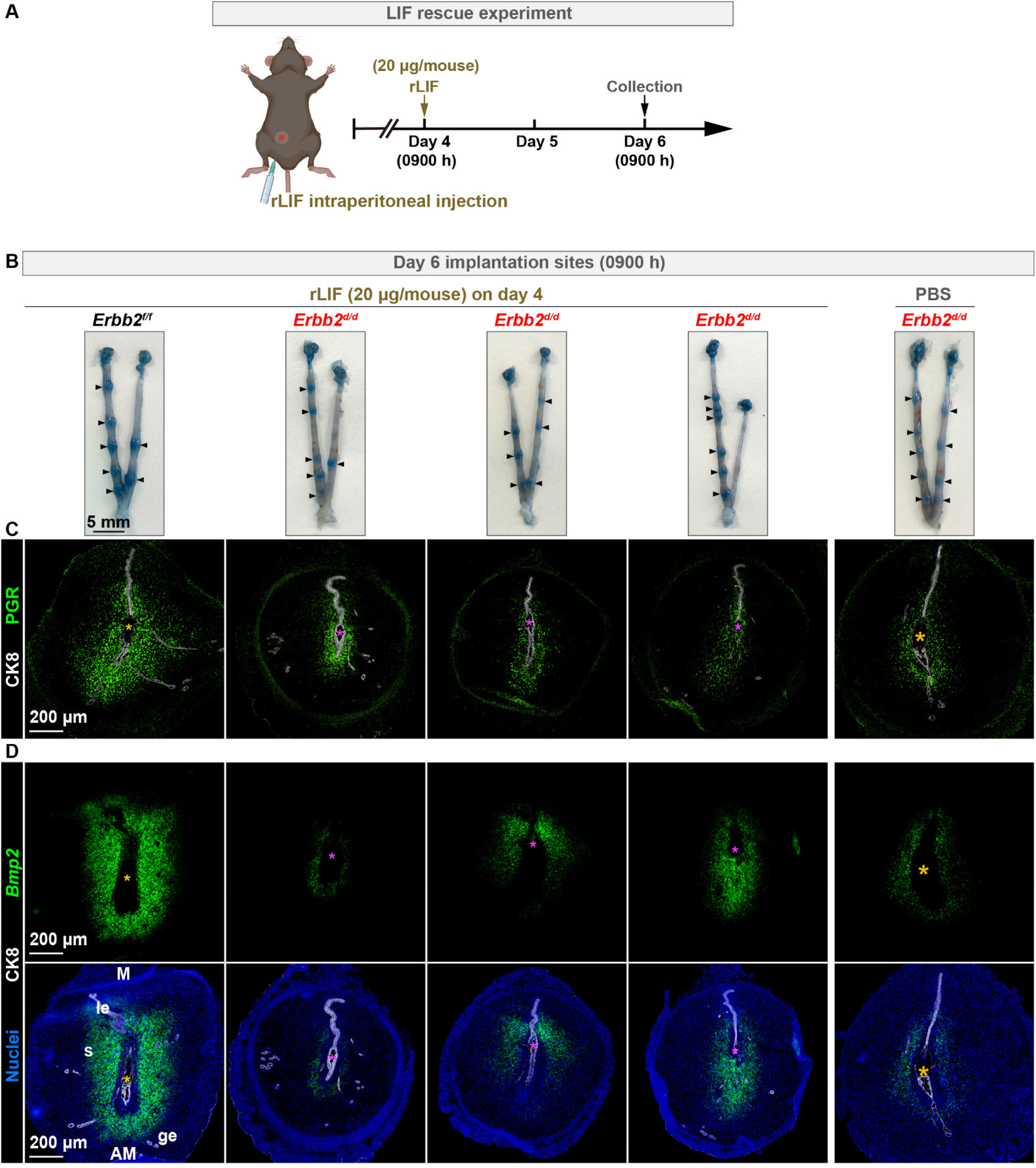
rLIF supplementation partially rescue embryo implantation in *Erbb2^d/d^* mice. (**A**) Schema of experimental design. Recombinant LIF (rLIF, 20 µg/mouse) was intraperitoneally injected into *Erbb2^f/f^* and *Erbb2^d/d^* mice at 0900 h on day 4 of pregnancy, and uteri were collected at 0900 h on day 6 of pregnancy (*Erbb2^f/f^* with LIF treatment: n = 3; *Erbb2^d/d^* with LIF treatment: n = 3; *Erbb2^d/d^* with PBS treatment: n = 3). (**B**) Representative images of implantation sites from *Erbb2^f/f^* and *Erbb2^d/d^* mice on day 6. Scale bar, 5 mm. (**C**) Immunofluorescence staining showing PGR and CK8 expression at implantation sites. Scale bar, 200 µm. (**D**) FISH of decidualization marker *Bmp2* and CK8. Scale bars, 200 µm.

**Supplemental Figure 7.**
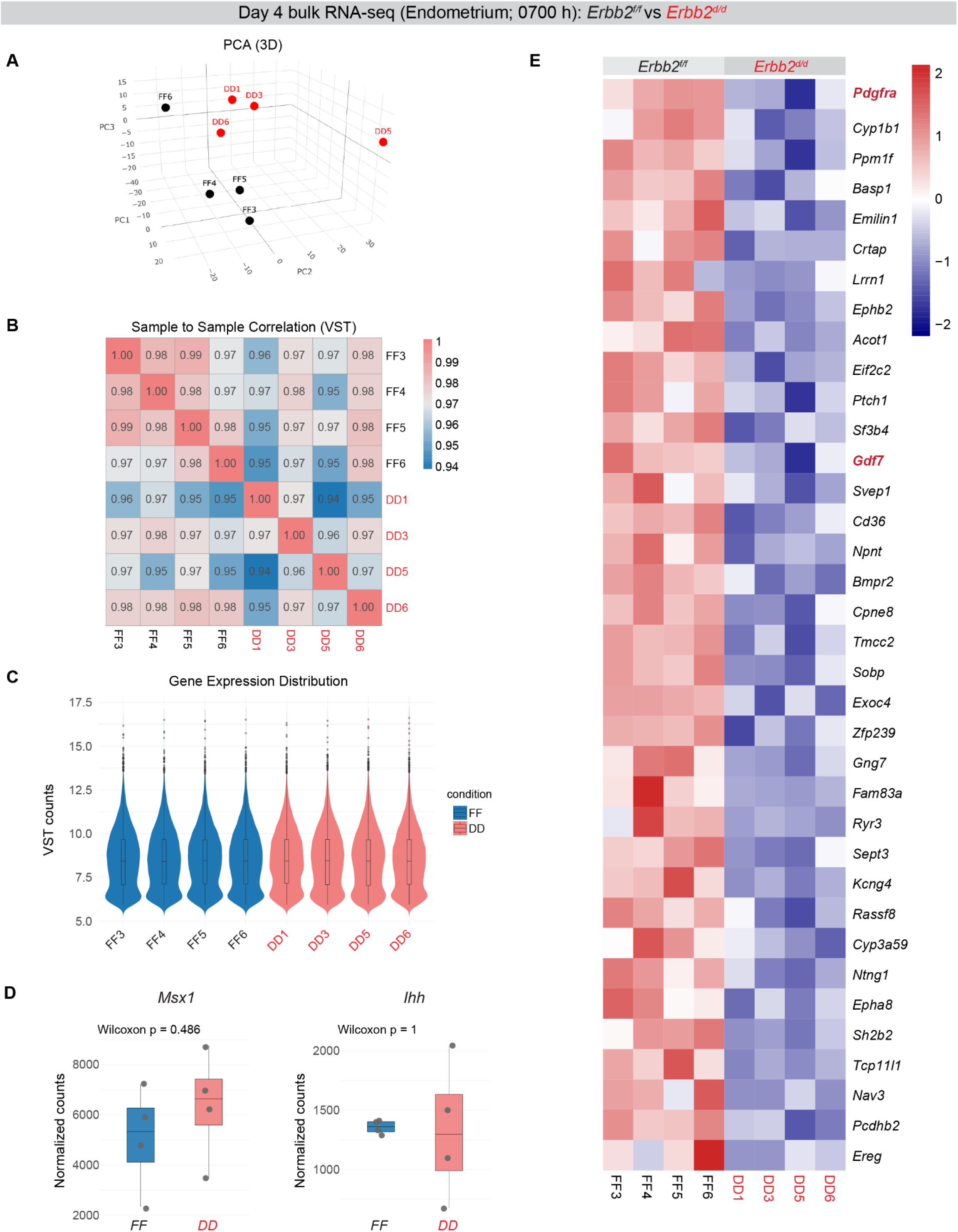
Bulk RNA-seq analysis of day 4 endometrium (0700 h) reveals stromal gene alterations in *Erbb2^d/d^* uteri. (**A**) 3D principal component analysis (PCA) of bulk RNA-seq data from day 4 (0700 h) endometrial tissues of *Erbb2^f/f^* (n =4) and *Erbb2^d/d^* (n =4) mice. (**B**) Sample-to-sample correlation heatmap based on variance-stabilizing-transformed (VST) counts. (**C**) Gene expression distribution (VST counts) across biological replicates. (**D**) Normalized expression of classical uterine receptivity markers, *Msx1* and *Ihh* (Wilcoxon test). (**E**) Heatmap of significantly downregulated genes in DD samples. Color scale indicates z-score– normalized expression.

**Supplemental Figure 8.**
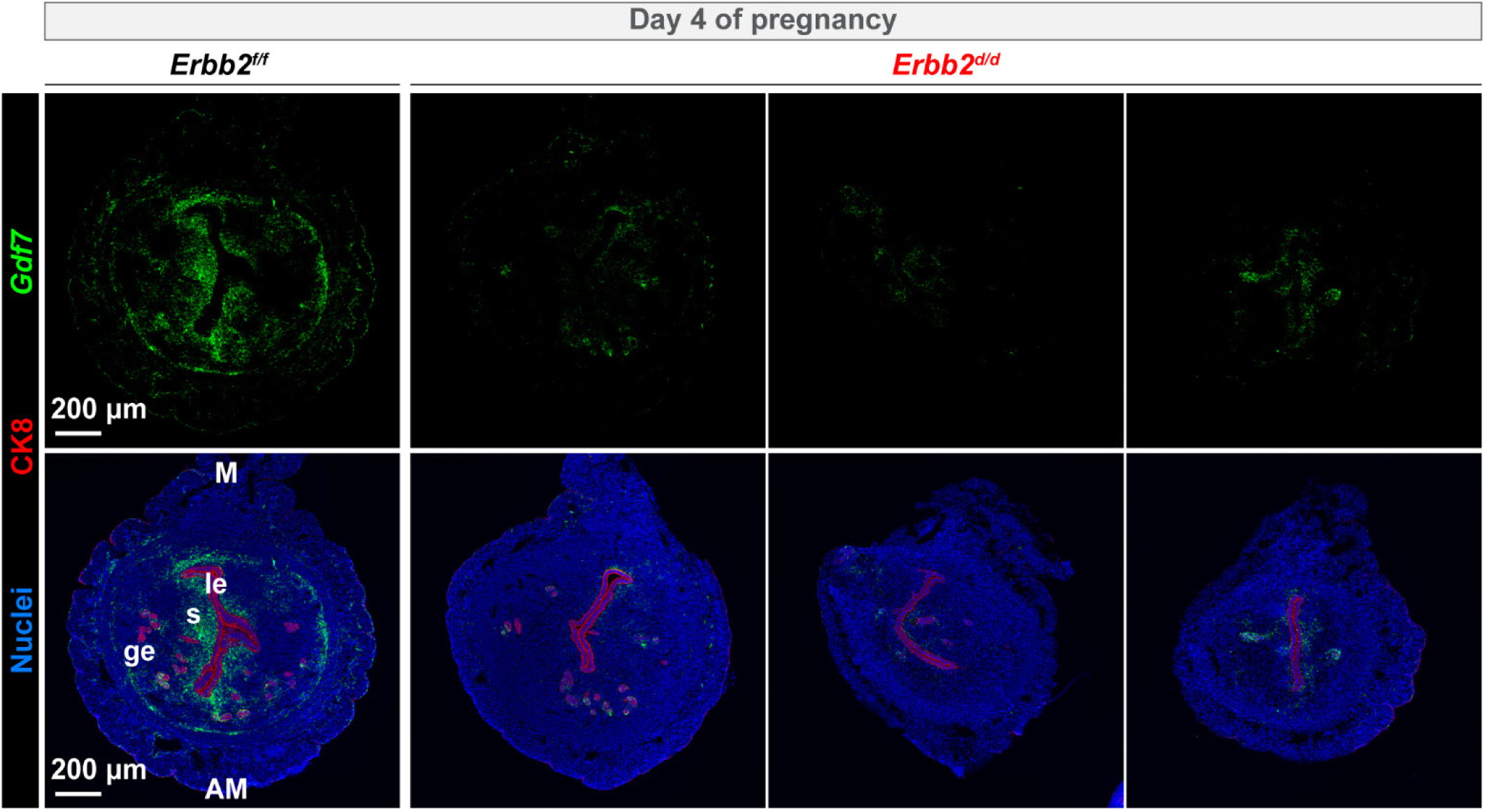
Stromal *Gdf7* was reduced in *Erbb2^d/d^* mice on day 4 of pregnancy. FISH of *Gdf7* and immunofluorescence of CK8 in *Erbb2^f/f^* (n = 3) and *Erbb2^d/d^* (n = 3) uteri on day 4 of pregnancy. Scale bar, 200 μm. le luminal epithelium, ge glandular epithelium, s stroma, M mesometrial pole, AM antimesometrial pole.

**Supplemental Table 1.** Clinical characteristics of women included in normal and RSA samples.

|  | normal (n=21) | RSA (n=21) | *P* |
| --- | --- | --- | --- |
| Age (years) | 26.71±2.51 | 27.86±3.24 | 0.209 |
| Gestation age (weeks) | 8.46±0.85 | 8.3±0.76 | 0.521 |
| BMI | 21.64±1.51 | 22.17±0.99 | 0.171 |
| smoking status | None | None |  |
| Previous unexplained pregnancy loss | 0 | 2-4 | < 0.0001 |
| Parity | 0-2 | 0-1 | 0.338 |
| Chromosomal analysis | - | normal | - |

**Supplemental Table 2.**
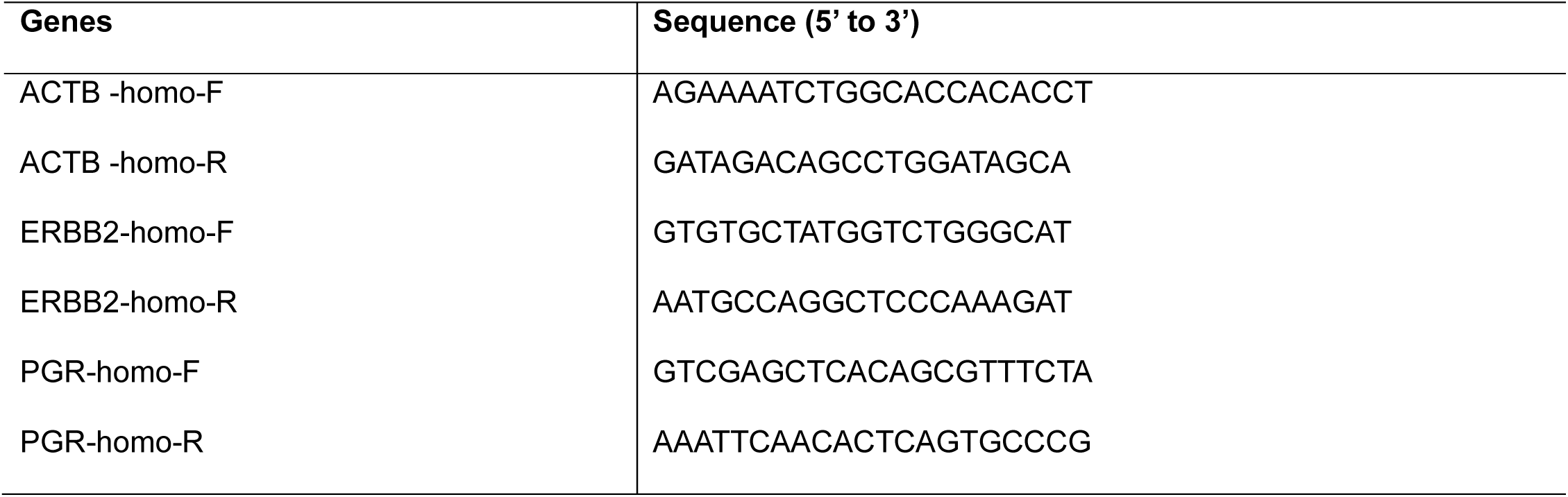
Sequences of primers used for qRT-PCR analyses.

